# Zika and Dengue Viruses Differentially Modulate Host mRNA Processing Factors Defining Its Virulence

**DOI:** 10.1101/2024.10.18.619084

**Authors:** Aaron Scholl, Binsheng Gong, Bingjie Li, Tahira Fatima, Nikki Tirrell, Spyros Karaiskos, Maria Rios, Joshua Xu, Sandip De

## Abstract

Rising global temperatures coupled with increasing international travel, and trade are contributing to spread of vectors such as ticks and mosquitoes, resulting in a surge of vector-borne flavivirus infection in human population. Furthermore, this increase in flavivirus infection pose threat to the safety of biologics such as cell and gene therapy products as human-derived materials are commonly used during manufacturing of these drug products. In this study, we conducted time-course transcriptomic and protein analyses to uncover the host molecular factors driving the virulence of Zika virus (ZIKV) and Dengue virus (DENV) in the context of host defense mechanisms, as these two viruses have caused the most recent and significant flavivirus outbreaks. Compared to DENV, ZIKV exhibited stronger virulence and cytopathic effects. RNA sequencing analysis revealed differential expression of various cellular factors, including RNA processing factors. Further investigation identified cell-type and time-dependent upregulation nonsense-mediated RNA decay (NMD), RNA degradation factors and nuclear pore complex (NPC) transcripts. Protein analysis showed that ZIKV, unlike DENV, degrades NMD factors in host cells, which along with mis-regulation of RNA degradation factors resulted in accumulation in host intronic transcripts as revealed by RNA-seq data. We also found that active nuclear transport is required for ZIKV replication, suggesting that therapeutic targeting of the NPC could potentially be effective in controlling ZIKV infection. Furthermore, from our findings we hypothesize that, ZIKV, but not DENV, drives early host cell cytopathy through targeted protein degradation. Current studies are underway to develop novel strategies to detect ZIKV, DENV and other flaviviruses in biologics based on transcriptomics and proteomics.

**Teaser:** Exploring the molecular basis of flavivirus virulence in host cells.

## Introduction

Although over 70 different flaviviruses have been identified, not all are known to cause disease in humans. While flaviviruses are predominantly transmitted to humans through vectors like mosquitoes and ticks, additional modes of transmission have been documented, such as sexual and vertical routes (*1, 2*), and through human donor tissues (*3*). Humans with flavivirus infection exhibit a wide range of symptoms, from mild skin rashes and flu-like symptoms (*4, 5*) to more severe outcomes like hepatomegaly (*4*), multi-organ failure (*6*), meningoencephalitis (*7*), and even death (*8, 9*). Moreover, infections during pregnancy can result in microcephaly and other severe developmental issues in the fetus (*10*). These studies highlight the wide range of tissue tropism and virulence observed among flaviviruses.

While flaviviruses are generally not linked to chronic infections, evidence has shown that these viruses can be present in specific cells for prolonged durations, even after clinical symptoms have subsided. Persistent infection by tick- and mosquito-borne flaviviruses has been documented in several studies, emphasizing the potential of these viruses to maintain a presence in human cells long after the initial infection (*11–13*). For example, ZIKV has been shown to persist in infected individuals for many months (*14*), and DENV-3 has been reported to establish persistent infection, particularly in immunocompromised patients (*15*). Flavivirus persistence can lead to long-lasting transcriptomic changes within the host cells even after the viral clearance (*16*), and long-term consequences of flavivirus infections in healthy adult hosts have also been reported (*17*).

In this study, we studied the mechanisms that may contribute for short- and long-term host defense against ZIKV and DENV infections, which exhibit differing virulence. Our data revealed distinct transcriptomic and protein profiles in host cells following ZIKV and DENV infection. We found differential effect of ZIKV and DENV on various cellular factors, which may contribute to dysregulation of several cellular pathways and promote virulence. Interestingly, we also found upregulation of transcripts of several host factors involved in mRNA processing, such as RNA degradation factors, splicing factors and nonsense-mediated RNA decay (NMD) factors, differed in a cell type-specific manner. Notably, while the master regulator of the NMD pathway, *UPF1*, showed increased transcript levels by both ZIKV and DENV, it was selectively degraded at the protein level by ZIKV but not by DENV-3. Similarly, expression of *UPF2*, another co-factor in the NMD pathway, was reduced at both the RNA and protein levels during ZIKV infection but not DENV-3. ZIKV infection also resulted in the accumulation of intronic RNA transcripts, potentially due to disruptions in the NMD and RNA degradation pathways. This was also observed following long-term infection (>3 months) of cells with DENV-3 infection but not during acute infection (≤5 days). We also found cell type-specific upregulation of nuclear pore complex (NPC) factors upon ZIKV and DENV infection, suggesting important role of NPC during ZIKV and DENV infection. Consistent with this, pre-treatment with Leptomycin B (LMB), an inhibitor of nuclear export, reduced ZIKV infection, underscoring the NPC’s pivotal role in promoting ZIKV infection and its potential as a target for novel antiviral therapies.

## Results

### Vero cells are less susceptible to DENV-3 compared to ZIKV

Vero E6 were infected with ZIKV (MR766 strain) or DENV (serotype 3) at a multiplicity of infection of 1 (MOI-1), and harvested at 1-, 3-, and 5-days post infection (DPI). Cells were analyzed for expression of viral envelope protein (env, ZIKV) or non-structural protein 1 (ns1, DENV) by western blot and immunocytochemistry (ICC) analysis and for viral RNA expression by RNA sequencing (RNA-seq). ZIKV env protein was detected as early as 1-DPI; however, DENV ns1 was only detected at 5-DPI (**Fig. 1A, B**). Consistent with this, RNA-seq analysis detected high levels of ZIKV transcripts in Vero cells at 3- and 5-DPI, but DENV transcripts were only detected at 5-DPI and were significantly low than ZIKV (**Fig. 1C**).

**Fig. 1.**
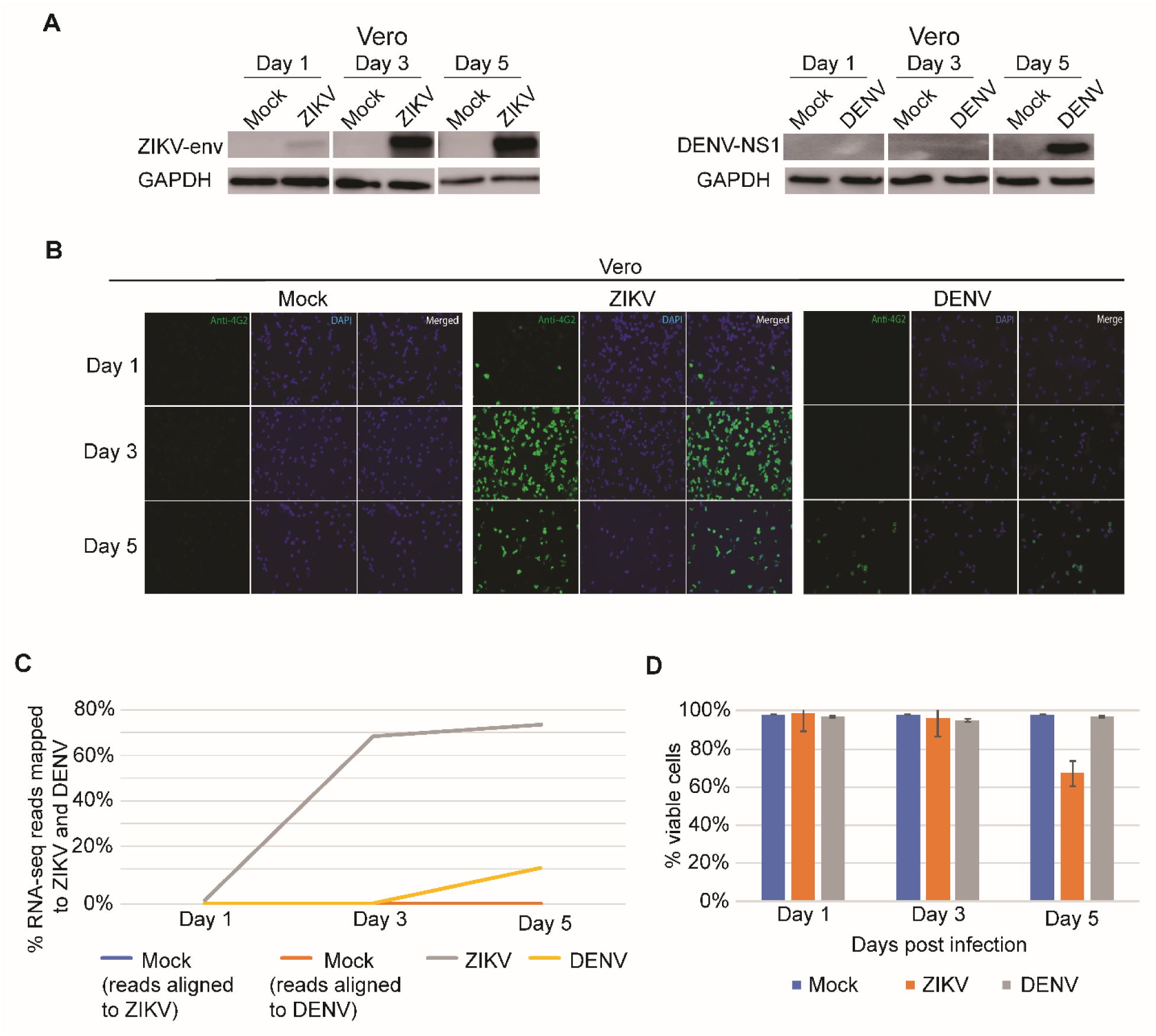
Vero cells are less susceptible to DENV-3 compared to ZIKV-MR766. (A) Western blot analysis of Vero infected with ZIKV (left) and DENV (right) at 1-, 3-, and 5-DPI probed with anti-ZIKV envelope, anti-DENV-NS1, respectively and anti-GAPDH. (B) Immunocytochemistry assay of mock-infected control, ZIKV-, and DENV-infected Vero cells at 1-, 3-, and 5-DPI. Antibodies used were anti-4G2 (anti-flavivirus, green) and DAPI (blue). (C) Percent of total RNA-seq reads that mapped to ZIKV-MR766 (grey-infected Vero and blue-mock Vero samples) and to DENV-3 (yellow-infected Vero and orange-mock Vero samples) at 1-, 3-, and 5-DPI. (D) Percent cell viability for mock-infected control (blue), Vero ZIKV (orange) and Vero DENV (grey) samples on 1-, 3-, and 5-DPI.

We also investigated the cytopathic effect (CPE) in Vero cells upon ZIKV and DENV infection. There was no significant difference in cellular viability between mock-infected control and ZIKV-infected cells until 5-DPI in Vero cells; however, DENV infection did not cause any Vero cell death up to 5-days (**Fig. 1D**). Together, these data suggest that Vero cells are highly susceptible to ZIKV compared to DENV.

### RNA sequencing of ZIKV and DENV infected cells, and pathway analysis

To understand the molecular basis of ZIKV and DENV virulence with respect to host defense mechanisms, we carried out RNA sequencing (RNA-Seq) of Vero cells following acute (days 1, 3 and 5) infection. Additionally, we carried out RNA-seq in Vero cells following long-term infection (LTI) with ZIKV or DENV (**Fig. S1)**, Principal component analysis (PCA) of RNA-Seq data from Vero cells infected with ZIKV or DENV demonstrate the close association between the samples within the test condition (**Fig. 2A**). For example, the mock-infected Vero cell samples at 1 day post infection (D1-Cont) are closely associated with each other and are distinct from ZIKV-infected samples (**Fig. 2A**). As ZIKV infection progresses to 3- and 5-days post infection (DPI), the infected samples increasingly diverge from the mock-infected controls, and the separation becomes pronounced in the LTI samples (**Fig. 2A**). Interestingly, the DENV-infected samples following acute infection are more closely associated with the respective mock-infected controls for each time point (**Fig. 2A**). For DENV, only LTI samples were clearly separated from mock-infected controls (**Fig. 2A**).

**Fig. 2.**
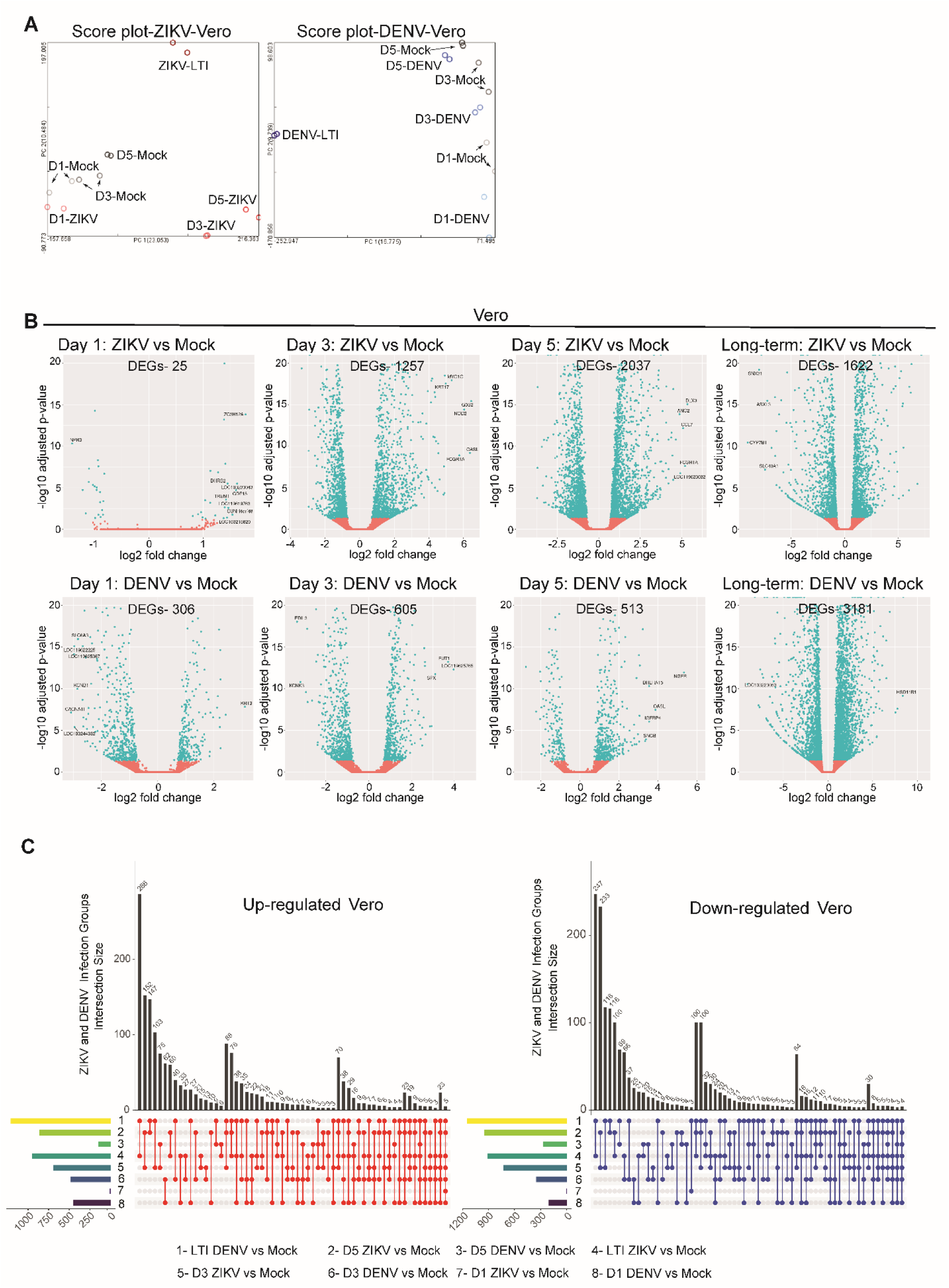
Analysis of differences between infected and mock samples using RNA-seq. (A) Principal component analysis (PCA) of mock (Cont), ZIKV infected Vero, and DENV infected Vero samples for 1-, 3-, and 5-DPI and for long-term infection samples used in RNA-seq analyses. Two replicates are graphed per condition per time point. (B) Volcano plots of −log_10_ adjusted p-value versus log_2_ fold change of differentially expressed genes between mock versus ZIKV- or DENV-infected samples at 1-, 3-, and 5-DPI, as well as mock versus long-term infected ZIKV or DENV samples. DEGs with adjusted p-value >0.05 are shown in red while DEGs with adjusted p-value <0.05 are shown in teal. (C) UpSet plots of up-regulated and down-regulated genes shared between mock, ZIKV-, and DENV-infected Vero sample datasets.

Volcano plots were generated based on differentially expressed genes (DEGs) for virus-infected Vero cells at each time point compared to the mock-infected controls (**Fig. 2B**). At 1-DPI, a low number of DEGs was observed in Vero cells following ZIKV infection; however, as the infection progressed the DEGs increased for each time point with the largest number of DEGs observed at 5-DPI (**Fig. 2B**). Long-term infection with ZIKV also resulted in many DEGs (**Fig. 2B**). Interestingly, this trend was not observed for DENV infection as compared to ZIKV, higher number of DEGs were observed following DENV infection at 1-DPI (**Fig. 2B**). For DENV infection, DEGs did not increase to the same extent as observed with ZIKV infection with slight decrease observed at 5-DPI compared to 3-DPI (**Fig. 2B**). Unlike, long-term infection with ZIKV, long-term DENV infection resulted in the highest number of DEGs (**Fig. 2B**).

Next, UpSet plots were generated for Vero ZIKV and Vero DENV samples to show intersections of DEGs between different comparison groups (**Fig. 2C**). Interestingly, long-term DENV and ZIKV samples not only exhibit the highest and second-highest number of DEGs, but they also share a greater number of DEGs compared to any other group. The next highest DEG pairing is between 3- and 5-DPI for ZIKV. This similarity between long-term infected cell lines suggests that mechanisms required for establishing long term infection *in vitro* may be shared between these two flaviviruses.

Gene set enrichment analysis (GSEA) was performed for ZIKV- and DENV-infected Vero cells using RNA-Seq data. As expected, several pathways were either up- or down-regulated following virus infection (**Fig. S2**). The analysis revealed that on 1-DPI, RNA processing pathways were suppressed for Vero ZIKV-infected samples compared to the respective mock-infected controls (**Fig. S2A**). At 3-DPI, the cells responded to the presence of ZIKV via activation of immune response, defense response, and response to external stimulus pathways, while cell division-related processes such as chromosome organization and segregation were suppressed. By 5-DPI, the virus was suppressing ribosome-related pathways while immune-related pathways were activated. In long-term ZIKV infection samples, the cell division-related pathways were reactivated. Consistent with previous studies demonstrating suppression of ciliogenesis and motile ciliary functions (*18*), we also observed cilium and cell-adhesion related pathways were suppressed in long-term ZIKV-infected Vero samples.

In DENV-infected Vero samples at 1-DPI, nuclear transport and nucleocytoplasmic transport pathways were activated while cilium-related pathways were suppressed (**Fig S2B**). At 3-DPI, cell division-related pathways were activated along with ncRNA metabolic processes while cilium-related pathways were suppressed. By 5-DPI, pathogen defense-associated pathways were activated, and cell cycle pathways were repressed. In long-term Vero DENV-infected samples, pathogen defense-associated pathways remain activated, and the cilium-related pathways remain suppressed.

To further expand on identifying host factors important for flaviviral defense, we infected SK-N-SH cells, a human neuroblastoma cell line, with ZIKV and carried out RNA-seq (acute infection 1-, 3- & 5-DPI samples, we could not derive long-term ZIKV-infected SK-N-SH cells). As with ZIKV-infected Vero samples, we investigated the CPE in SK-N-SH cells upon ZIKV infection. Difference in cellular viability between mock and ZIKV-infected cells was not obvious until 5-DPI (**Fig. S3A**). RNA-seq analysis detected high levels of ZIKV transcripts in SK-N-SH samples at 3- and 5-DPI, similar to the trend observed in ZIKV-infected Vero samples (**Fig. S3B**). PCA analysis for SK-N-SH mock and ZIKV-infected samples showed that there is less of a difference on 1-DPI while 3- and 5-DPI show a greater level of divergence nearly mimicking the trend of Vero mock vs ZIKV-infected samples (**Fig. S4A**). In the volcano plots for ZIKV-infected SK-N-SH samples, starting at 1 DPI, there are very few DEGs, similar to the volcano plot for ZIKV-infected Vero at 1-DPI (**Fig. S4B**). ZIKV-infected SK-N-SH at 3- and 5-DPI shared an identical trend of increased DEGs compared to ZIKV-infected Vero of the respective time points as the infection grows (**Fig. S4B**). UpSet plots were generated for SK-N-SH ZIKV-infected samples to show intersections of DEGs between different comparison groups (**Fig. S4C**). As with Vero UpSet plots, ZIKV-infected samples from 3- and 5-DPI shared the highest number of shared DEGs. GSEA data for ZIKV-infected SK-N-SH at 1-DPI showed most of the top activated pathways being related to transforming growth factor beta (TGF-β) signaling. TGF-β is known to regulate the inflammatory immune response, so the presence of this pathway in our GSEA dataset for early infection samples is expected (*19, 20*). However, by 3- and 5-DPI, mRNA processing-related pathways were activated in addition to ribosome and ribogenesis-related pathways (**Fig. S4D**). At both 3- and 5-DPI, cell adhesion-related pathways were suppressed. It has been recently reported that ZIKV interacts with various aspects of actin and tubulin cytoskeleton reorganization (*21–23*). While cytoskeletal reorganization plays its own part in flaviviral replication, it may also be another aspect of the overall subversion of the circadian clock as a means for the virus to gain control of the host cell (*24, 25*). Changes to the circadian clock during flavivirus infection have been identified in several recent studies (*26, 27*). The similarities between GSEA results for Vero and SK-N-SH ZIKV infections led us to delve deeper into mRNA processing pathways.

Given mRNA processing and RNA splicing pathways were differentially expressed in one Vero and two SK-N-SH samples and previous proteomic and siRNA studies have shown flavivirus infection regulation in host cells via RNA processing factors (*28–30*), we searched for mRNA processing and other related GO annotations in the GSEA dataset. GSEA data was significant for mRNA processing (GO:0006397), mRNA metabolic process (GO:0016071), and RNA splicing (GO:0008380) pathways at 1-DPI for Vero ZIKV along with mRNA processing, mRNA metabolic process, and RNA splicing at 1-, 3- and 5-DPI for Vero DENV compared to respective samples (**Table S1**). Despite the significance found in these pathways, the GSEA dot plots only cover the 10 most significant activated and repressed pathways of each dataset, which excludes mRNA processing from GSEA figures (**Fig. S2 & S4**).

While GSEA considers all genes within the dataset to determine enriched pathways, overrepresentation analysis (ORA) uses only the differentially expressed genes within the dataset. In the ORA results between mock and ZIKV- or DENV-infected datasets, actin cytoskeleton and cell junction pathways emerge (**Fig. S5**). Regarding the top 20 pathways in the ORA results of ZIKV-infected SK-N-SH, the only pathway not related to development or differentiation at 1-DPI was viral genome replication pathway, which is to be expected with the limited number of DEGs at this time point (**Fig. S6**). At 3-DPI, however, two main pathway groupings appear: response to endoplasmic reticulum (ER) stress and the transmembrane receptor protein serine/threonine kinase signaling pathway. Members of this signaling pathway participate in MAPK signaling pathway, TGF-β signaling pathway, and adherens junctions. The TGF-β signaling pathway has been reported to interact with the molecular circadian oscillator, amongst other functions such as regulating immune response (*20, 31*). The pathways related to ER stress are to be expected with flaviviral replication occurring in the ER (*32, 33*). Furthermore, several pathways related to unfolded and topologically incorrect proteins emerge. By 5-DPI, the main two groups of pathways were RNA splicing- and cell cycle-related. RNA splicing and usurping ribosomal control have been reported in many families of viruses as a means of taking over host post-transcriptional regulation (*34, 35*). Cell cycle dysregulation is a direct consequence of RNA splicing pathway alterations. As various RNA processing and cellular response to unfolded/topologically incorrect proteins pathways appeared in the ORA dot plots in addition to identification of mRNA processing and related pathways in GSEA analysis, we next studied the transcript dynamics of NMD pathway factors due to its function in RNA quality control.

### Nonsense mediated decay pathway transcripts and proteins are mis-regulated upon flavivirus infection

A heatmap was generated to analyze the dynamics of NMD pathway transcripts across various infection time points (**Fig. 3A**). The heatmap from Vero samples showed increased *UPF1* transcript levels across all time points of acute and long-term infection (**Fig. 3A**). This result is supported by normalized transcript-reads per million when looking at *UPF1* via Integrative Genomics Viewer (IGV) data (**Fig. 3B**). However, at the protein level, we observed a gradual decrease in UPF1 levels during acute ZIKV infection, but an increase during acute DENV infection in respective western blots (**Fig. 3C**). A similar result is observed in immunocytochemistry assays for UPF1 in ZIKV- and DENV-infected Vero samples (**Fig. 3D**). It is reported that the ZIKV capsid protein targets UPF1 for degradation (*36*). Despite increased *UPF1* transcripts for ZIKV and DENV during acute infection, UPF1 proteins in ZIKV-infected samples decrease over the same timeframe. This finding suggests that ZIKV-MR766, but not DENV-3, can degrade UPF1 proteins during acute infection. Despite sequence similarity of 55.6% between ZIKV and DENV, UPF1-targeting viral capsid proteins exhibit a high degree of similarity between flaviviruses (*37, 38*). These differences could account for functional changes in UPF1-degradation capabilities, limiting the overall subversion of NMD in DENV-infected samples, but not in ZIKV-infected samples. An interesting trend emerged when comparing the NMD heatmaps of ZIKV-infected Vero transcripts to DENV-infected Vero transcripts. ZIKV-infected Vero samples see a reduction in more than half of the transcripts at 1- (13/20 decreased) and 3-DPI (13/20 decreased), while 5-DPI (8/20 decreased) and long-term infection (4/20 decreased) there are increased transcripts for more than half of the genes (**Fig. 3A**). Interestingly, the opposite trend is present in DENV-infected Vero samples. At 1- (0/20 decreased) and 3-DPI (3/20 decreased), nearly all NMD-related gene transcripts exhibit increased levels, while there is a general decrease in transcript levels at 5-DPI (12/20 decreased) and an even distribution of increased and decreased transcripts during long-term infection (9/20 decreased) (**Fig. 3A**).

**Fig. 3.**
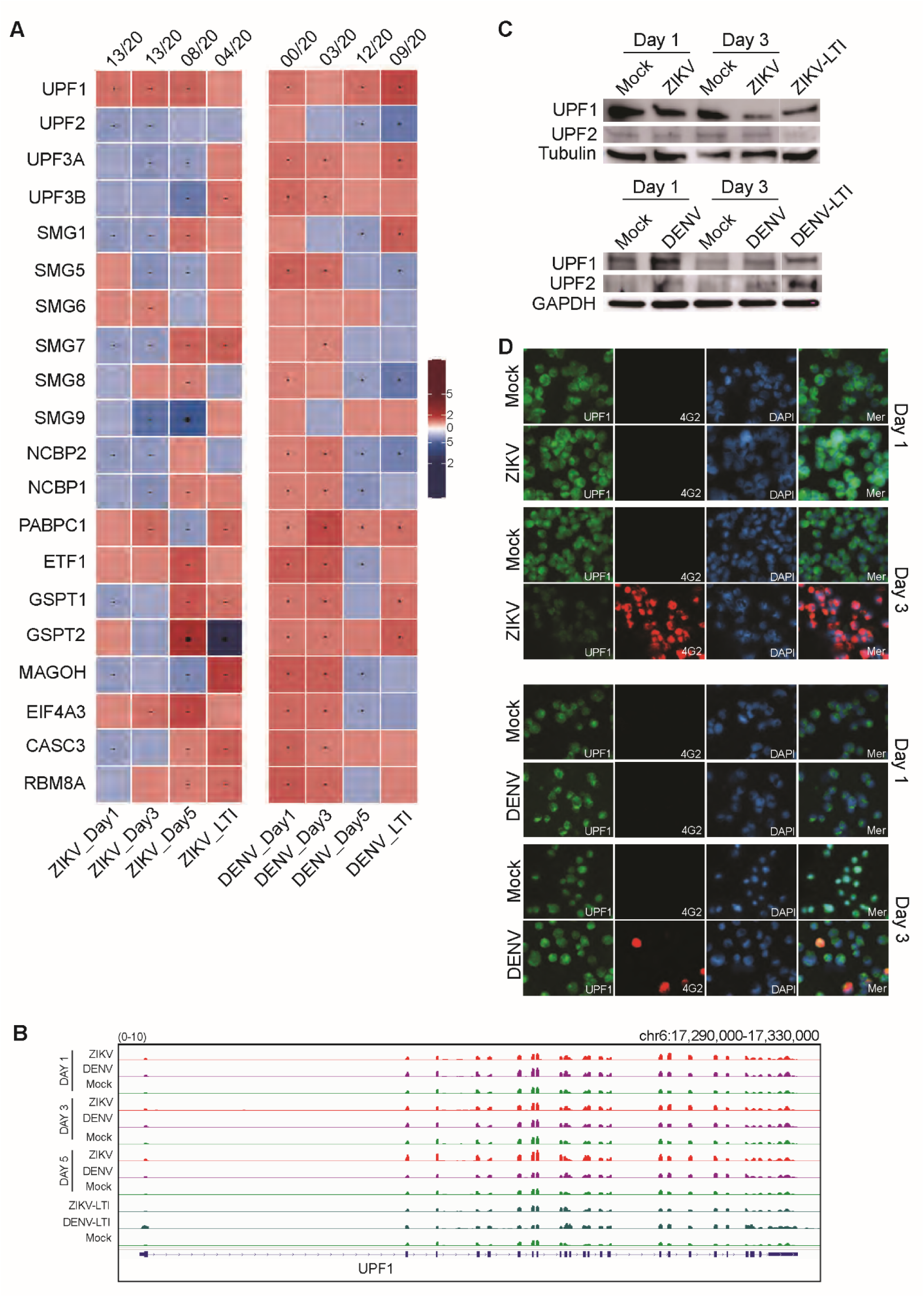
The up-regulation of NMD genes reveals distinct differences among ZIKV and DENV. (A) Heatmap of normalized RNA-seq Z-score data for mock versus ZIKV-infected (left) and DENV-infected (right) Vero samples at 1-, 3-, and 5-DPI and long-term infected samples. (B) A comparison of normalized RNA-seq reads represented in reads per million for UPF1 and visualized in IGV. Comparisons shown for mock, ZIKV-, and DENV-infected Vero samples at 1-, 3-, 5-DPI, and long-term infected samples. (C) Western blot analysis of anti-UPF1, anti-UPF2, and anti-alpha tubulin or anti-GAPDH as housekeeping controls for Vero acute ZIKV and DENV infections at 1-, 3-DPI and long-term infections (LTI). (D) Immunocytochemistry staining of Vero mock, ZIKV, and DENV at 1- and 3-DPI. Antibodies used were anti-UPF1 (in green), anti-4G2 (anti-flavivirus, in red), and DAPI (in blue).

In ZIKV-infected human SK-N-SH samples, we observed increased transcripts for most NMD-related genes, with overall increased transcript levels at 1- (5/20 decreased), 3- (5/20 decreased), and 5-DPI (7/20 decreased) (**Fig. S7A**). UPF1 transcripts were decreased in ZIKV-infected SK-N-SH samples at 1-DPI but increased at 3- and 5-DPI. At the protein level, UPF1 in ZIKV- and DENV-infected SK-N-SH samples indicated different trends (**Fig. S7B, C**). UPF1 proteins decrease only at 5-DPI in ZIKV-infected SK-N-SH samples, but no notable changes between mock and DENV-infected SK-N-SH samples were noted at any time point.

At the transcript level, *UPF2* is reduced in ZIKV- and DENV-infected Vero cells for all time points except 1-DPI (**Fig. 3A**). However, at the protein level, there was a decrease in UPF2 expression in ZIKV-infected Vero cells but not in DENV-infected Vero cells (**Fig. 3C, S8A**). Interestingly, *UPF2* transcripts were increased at 1-, 3-, and 5-DPI for ZIKV-infected SK-N-SH (**Fig. S7A**), and there was a notable decrease in UPF2 at the protein level by 5-DPI (**Fig. S7B, S8A**). This difference could be due to degradation of UPF2 via the viral capsid. At the protein level, UPF2 exhibited no distinct changes between mock and infected samples for DENV-infected SK-N-SH (**Fig. S7B, S8A**). The transcript heatmap findings for UPF2 were visualized in IGV by normalized transcript-reads per million, IGV findings support our heatmaps from ZIKV- and DENV-infected Vero samples and ZIKV infected SK-N-SH samples (**Fig. 3B**).

Heatmaps were generated for NMD target genes for ZIKV- and DENV-infected Vero and ZIKV-infected SK-N-SH samples (**Fig. 4A, B**). Between ZIKV- and DENV-infected Vero, several transcripts had consistent up- or down-regulation at all time points (**Fig. 4A**). The transcripts that were upregulated for both datasets were *NFKBIB* (8/8 conditions), *MID1IP1* (8/8 conditions), *GADD45A* (8/8 conditions), *CARS2* (8/8 conditions), and *ASNS* (7/8 conditions). Only *TRIM32* (8/8 conditions) exhibited down-regulation at all time points for both datasets. NMD target genes for ZIKV-infected SK-N-SH had varying results, with many transcripts exhibiting consistent up- or down-regulation at all time points (**Fig. 4B**). The transcripts that were similarly upregulated when compared to the Vero datasets were *NFKBIB* (10/11 conditions), *MID1IP1* (10/11 conditions), *GADD45A* (10/11 conditions), and *ASNS* (10/11 conditions). IGV comparisons of *ASNS* and *GADD45A* in flavivirus-infected Vero and SK-N-SH datasets showed increased transcript levels compared to mock datasets at similar time points, reflecting observations made with heatmap data (**Fig. 4C**). Additional IGV comparisons of *MID1IP1*, *NFKBIB*, and *RGS2* had similar trends during ZIKV infection (**Fig. S9A-C**). Interestingly, in ZIKV-infected SK-N-SH, we observed an increase in intronic reads in IGV, as indicated by arrowheads (**Fig. S9A-C**). As intronic reads accumulated, we investigated other RNA degradation control pathways to determine if changes occurred during ZIKV and DENV infection.

**Fig. 4.**
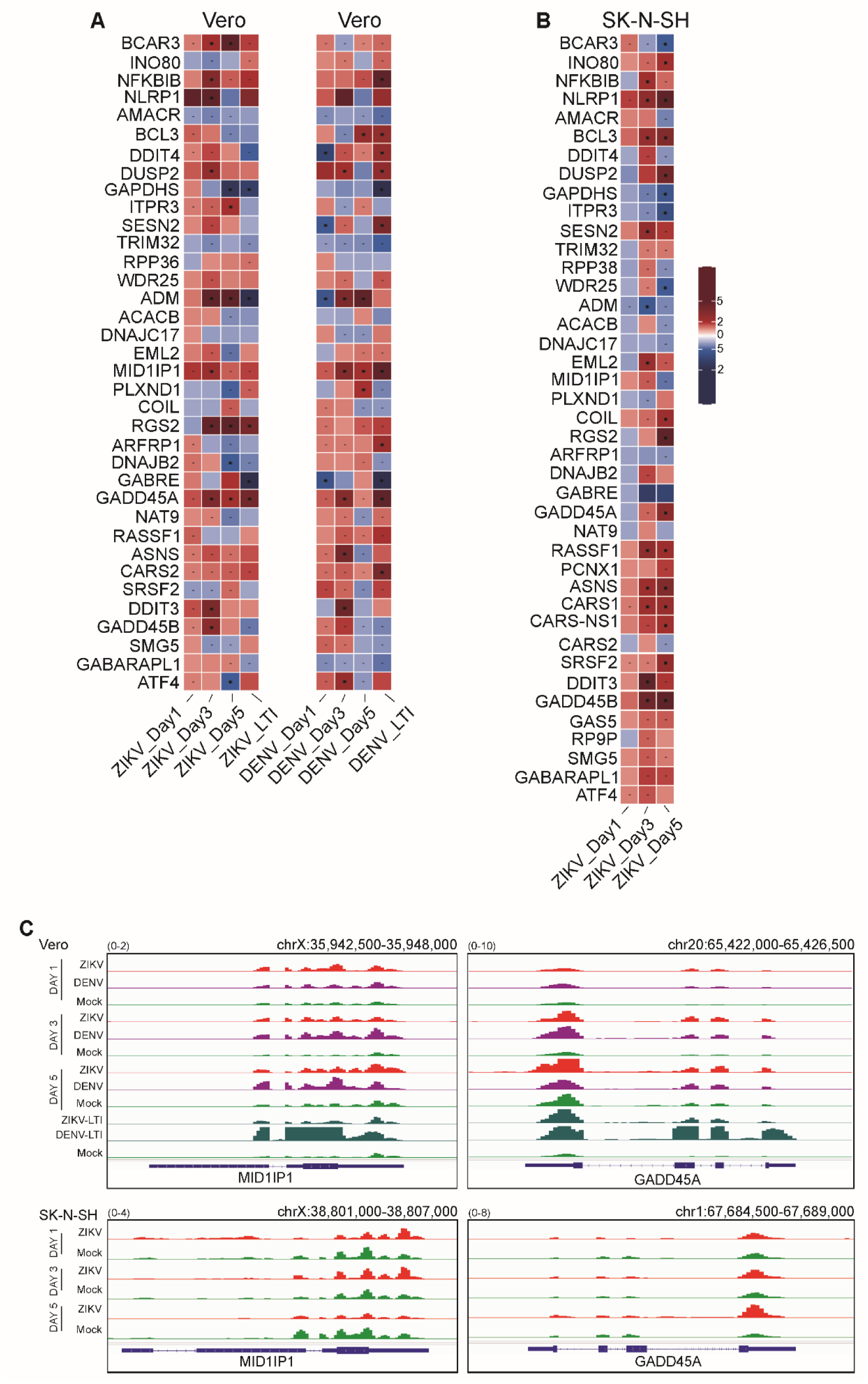
Comparison of NMD target transcripts in Vero, SK-N-SH acute and long-term infections. (A) Heatmap of normalized RNA-seq Z-score data for mock versus ZIKV-infected (left) and DENV-infected (right) Vero samples at 1-, 3-, 5-DPI, and long-term infected samples. (B) Heatmap of normalized RNA-seq Z-score data for mock versus ZIKV-infected SK-N-SH samples at 1-, 3-, and 5-DPI. (C) A comparison of normalized RNA-seq reads represented in reads per million for *MID1IP1* and *GADD45A* and visualized in IGV. Comparisons shown for mock, ZIKV-, and DENV-infected Vero and SK-N-SH samples at 1-, 3-, 5-DPI, and long-term infected samples.

### Analysis of RNA degradation pathway transcripts

Heatmaps were generated for RNA degradation-related genes for ZIKV- and DENV-infected Vero and ZIKV-infected SK-N-SH samples (*39*). As with NMD pathway heatmaps, RNA degradation pathway heatmaps had a general decrease in transcript levels for ZIKV-infected Vero samples at 1- (35/49 decreased) and 3-DPI (31/49 decreased) with a shift to increased transcript levels in more than half of the genes by 5-DPI (16/49 decreased) and long-term infection (20/49 decreased) (**Fig. 5A**). Similarly, in DENV-infected Vero samples, there was an increase in transcripts at 1- (8/49 decreased) and 3-DPI (11/49 decreased), but a major shift to decreased transcripts was apparent by 5-DPI (42/49 decreased) and long-term infection (26/49 decreased) (**Fig. 5A**). The heatmap from ZIKV-infected SK-N-SH samples indicated that transcripts were mostly upregulated upon infection (**Fig. S10A**). One transcript, *RNASEL*, was consistently increased at all time points in SK-N-SH ZIKV samples but decreased in Vero samples at all time points in both ZIKV and DENV infection (**Fig. S10A, Fig. 5A**). Normalized transcript-reads per million when viewed in IGV showed similar results to heatmap findings for *RNASEL* (**Fig. 5B**). We also checked the IGV profiles of *PAN2* and *LSM2* to confirm the heatmap findings from ZIKV-infected SK-N-SH samples and observed similar results (**Fig. S10B**). As the transcript level of RNA degradation factors are mis-regulated during flavivirus infection, we then compared the NGS read coverage (log2CPM) within 200 base pairs of intron/exon and exon/intron boundaries in both Vero mock, Vero ZIKV and Vero DENV (**Fig. 5C**) as well as in SK-N-SH mock and SK-N-SH ZIKV samples (**Fig. S10C**) to determine if there were any genome-wide accumulations of intronic sequences upon flavivirus infection. We observed that there were more genome-wide intronic reads present for ZIKV-infected Vero samples than Vero mock or DENV-infected Vero samples at 3- and 5-DPI. Interestingly, once ZIKV reach long-term infection, they show nearly similar coverage to mock. Intronic accumulation in some transcripts was also observed in Dengue infected cells but only after a long-term infection of DENV (**Fig. 5C, Fig. S9B-red arrowhead**). SK-N-SH ZIKV-infected samples had increased genome-wide levels of intronic reads by 5-DPI of infection over SK-N-SH mock (**Fig. S10C**). When UPF1 and UPF2 are degraded during acute ZIKV infection, they are unable to perform their main NMD-related function, and aberrant RNAs are not degraded. This further supports our findings that UPF1 and UPF2 are degraded during the progression of acute ZIKV infection. Accumulation of intronic signals was also observed in NMD target transcripts, RNA degradation pathway transcripts, nuclear pore complex pathway (NPC) transcripts, NMD pathway transcripts and interferon stimulated genes’ (ISGs) transcripts (**Fig. 5C, Fig. S10C**).

**Fig. 5.**
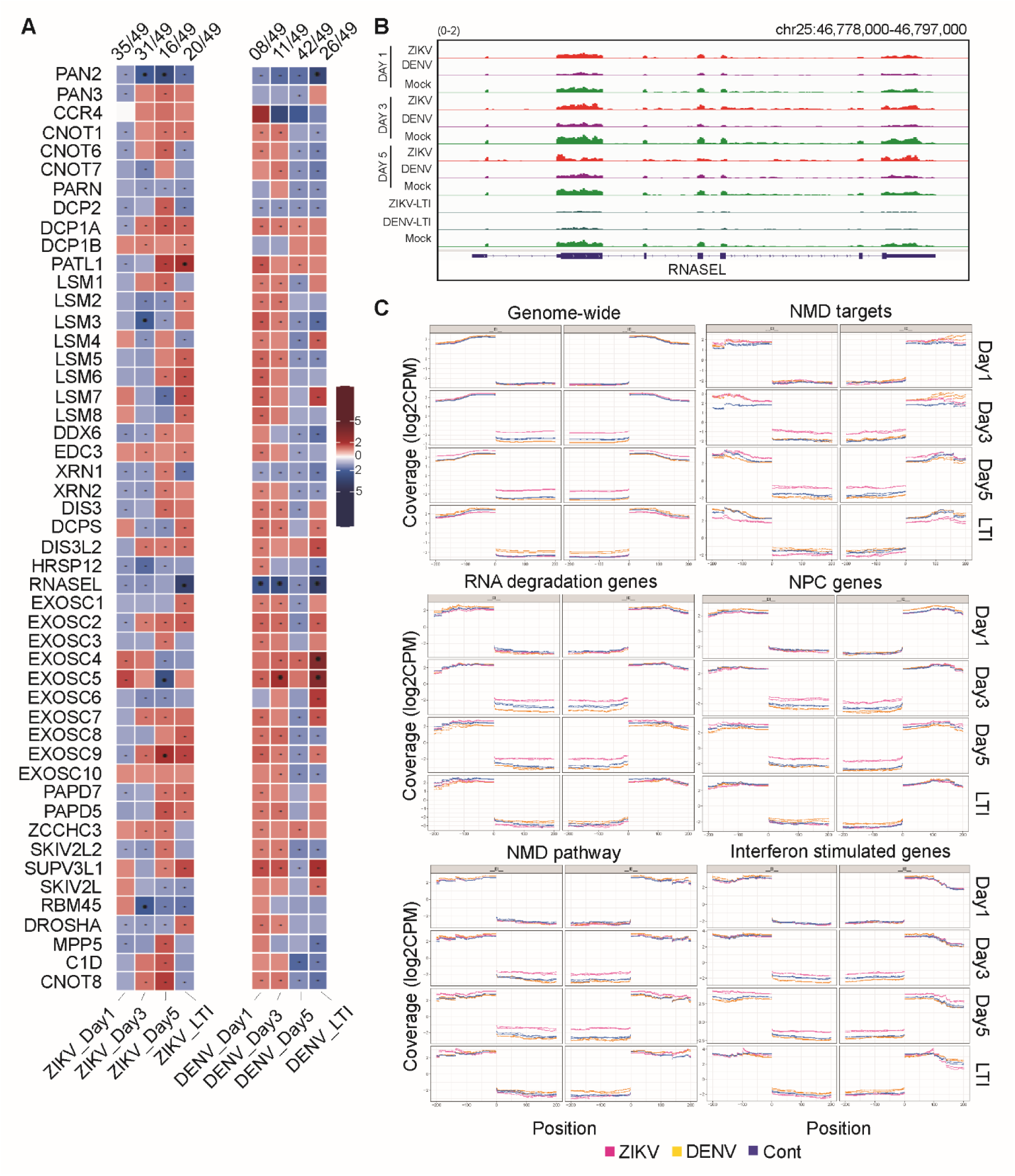
The up-regulation of RNA degradation pathway genes reveals distinct differences among ZIKV and DENV. (A) Heatmap of normalized RNA-seq Z-score data for mock versus ZIKV-infected (left) and DENV-infected (right) Vero samples at 1-, 3-, 5-DPI, and long-term infected samples. (B) A comparison of normalized RNA-seq reads represented in reads per million for RNASEL and visualized in IGV. Comparisons shown for mock, ZIKV-, and DENV-infected Vero samples at 1-, 3-, 5-DPI, and long-term infected samples. (C) Comparison of intron/exon and exon/intron boundaries genome-wide as well as for NMD pathway, NMD targets, RNA degradation genes, NPC genes, and ISGs within 200bp for mock (blue), ZIKV- (red), and DENV-infected (yellow) Vero samples.

### Nuclear pore complex transcripts mis-regulated upon flavivirus infection

Several flaviviral proteins exhibit nuclear localization and include nuclear export sequences (NESs) or nuclear localization sequences (NLSs), despite flaviviral replication occurring in the endoplasmic reticulum (*40, 41*). The nuclear pore complex (NPC) is also reported to be the target of flaviviral protease NS3 (*42*). Additionally, ZIKV NS5 protein is actively transported into the nucleus by NLS-mediated interactions with importin α and importin β1 heterodimer in the nuclear pore complex (*40*). Due to the reported nuclear localization of several flaviviral proteins, we investigated nuclear pore complex gene regulation.

Heatmaps were generated for NPC-related genes for ZIKV- and DENV-infected Vero and ZIKV-infected SK-N-SH samples. As with NMD and RNA degradation transcripts, Vero ZIKV samples were generally decreased in transcript levels at 1- (16/22 decreased) and 3-DPI (11/22 decreased) but had increased transcription by 5-DPI (9/22 decreased) and long-term infection (2/22 decreased) (**Fig. 6A**). The opposite trend is observed for DENV-infected samples. Samples exhibit an overall increase at 1- (2/22 decreased) and 3-DPI (4/22 decreased) but exhibit decreased transcript levels by 5-DPI (16/22 decreased) and a slight increase in overall transcript levels during long-term infection (9/22 decreased). We next investigated *NXT1*, a key factor in the nuclear export pathway, at both the transcript and protein level in Vero samples. *NXT1* had increased transcript and protein levels in Vero-ZIKV samples (**Fig. 6A, C, D**). Although in comparison to mock-infected control, *NXT1* is overexpressed at the transcript level, we did not see an increased level of NXT1 protein in Vero-DENV samples (**Fig. 6A, C, D**). Similarly, in ZIKV-infected SK-N-SH, despite increased transcript levels, no protein-level changes were observed (**Fig. S11A, C, D**). We also observed elevated intronic reads in ZIKV-infected Vero samples at 3- and 5-DPI (**Fig. S11B, black arrowhead**), but not in long-term ZIKV or in DENV-infected Vero samples (**Fig. 6B**). Accumulation of intronic reads was not observed in ZIKV-infected SK-N-SH samples at any time point post-infection (**Fig. S11B**).

**Fig. 6.**
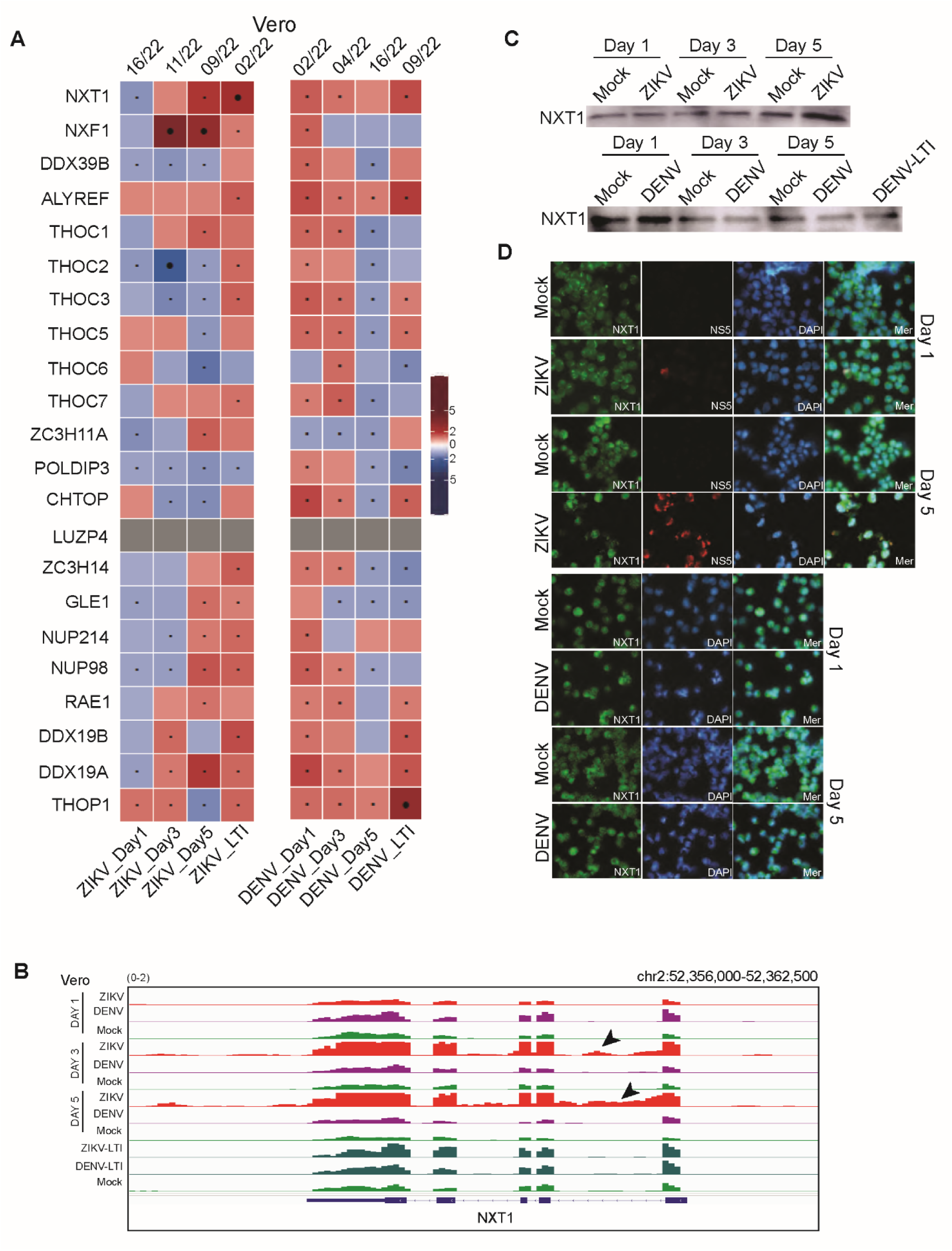
The up-regulation of NPC genes reveals distinct differences among ZIKV and DENV. (A) Heatmap of normalized RNA-seq Z-score data for mock versus ZIKV-infected (left) and DENV-infected (right) Vero samples at 1-, 3-, 5-DPI, and long-term infected samples. (B) A comparison of normalized RNA-seq reads represented in reads per million for NXT1 and visualized in IGV. Comparisons shown for mock, ZIKV-, and DENV-infected Vero samples at 1-, 3-, 5-DPI, and long-term infected samples. (C) Western blot analysis of anti-NXT1 for mock, ZIKV-, and DENV-infected samples at 1-, 3-, 5-DPI as well as long-term DENV. (D) Immunocytochemistry staining of mock, ZIKV, and DENV at 1- and 5-DPI. Antibodies used were anti-NXT1 (green), anti-ZIKV NS5 (red), and DAPI (blue).

Interferon-stimulated genes (ISGs) play a crucial role as the body’s first line of defense against viral infections (*43*) with intrinsically expressed ISGs playing a vital role in safeguarding stem cells (*44*). To explore whether this transcriptional pattern, as seen in the NMD pathway, RNA degradation complex, and NPC complex pathways, extends to ISGs in Vero-ZIKV and Vero-DENV samples, a heatmap was generated. Interestingly, we did not observe the expected up-regulation of ISG transcripts upon ZIKV and DENV infection in the tested cell lines (**Fig. S12**).

### Disruption of nuclear export via leptomycin B reduces ZIKV load in host cells

As previous studies have indicated interaction between flaviviruses and NPC components, and as we observed up-regulation of NPC transcripts in our study, we tested the role of the NPC complex in flavivirus infection. We inhibited nuclear export via treatment with 20nM leptomycin B (LMB) to determine if this would alter the infectivity of ZIKV (*22, 40, 42*). Cells were collected at 4-, 24-, and 48-hours post-infection (hpi) and subjected to ICC to observe changes to infectivity and cellular localization of viral and host components. Samples were immunoassayed with three anti-ZIKV antibodies, anti-4G2, anti-ZIKV NS5, and anti-ZIKV capsid, and the number of cells exhibiting signal was compared to the total number of cells (**Fig. 7A, B**, **Table S2**). In this experiment, two representative images were counted and averaged for each antibody at each respective time point. ZIKV-infected Vero cells untreated and treated with 20nM LMB treatment both showed 0% infection for all antibodies at 4hpi. By 24hpi, there were very minor differences in positive signal between untreated (6.09%, 4.12%) compared to treated (6.96%, 4.26%) for anti-4G2 and anti-ZIKV NS5, respectively. By 24hpi, positive signal for anti-ZIKV capsid antibody showed 54.93% infected cells for untreated and 42.17% infected cells for treated samples. At 48hpi, the results for all antibodies showed similar differences between untreated (71.77%, 67.29%, 77.34% positive signal) and treated (61.73%, 57.21%, 65.86% positive signal) samples for anti-4G2, -ZIKV NS5, and -ZIKV capsid, respectively (**Fig. 7A**). No CPE or morphological changes were observed upon 20nM LMB treatment in ZIKV-infected Vero cells.

**Fig. 7.**
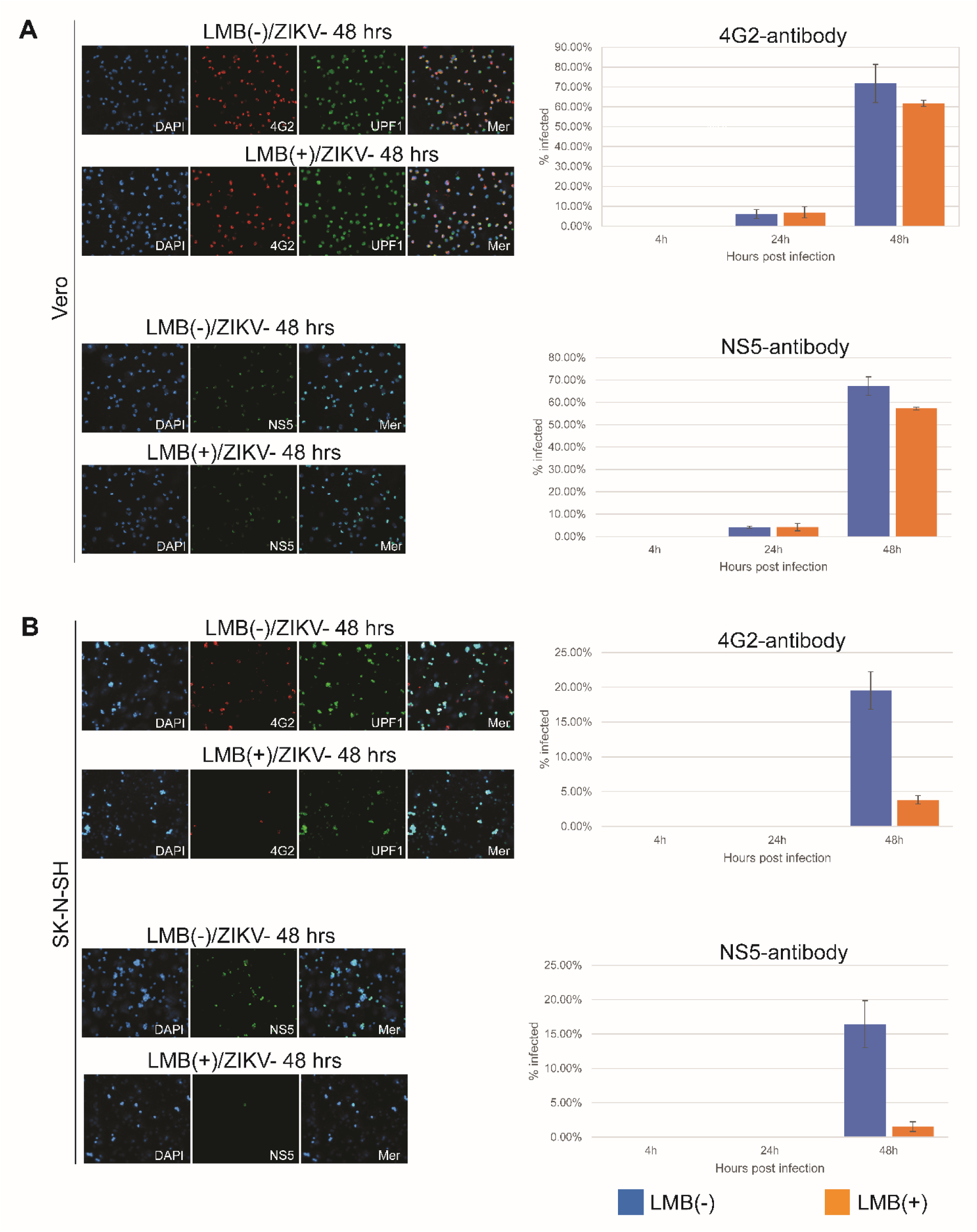
Disruption of nuclear export via leptomycin B (LMB) reduces ZIKV load in host cells. (A) Top panel-Immunocytochemistry of LMB-treated and untreated Vero samples at 48 hours. Antibodies used were anti-UPF1 (in green), anti-4G2 (in red), and DAPI (in blue). Shown on the right is a graphical representation of the percentage of infected cells at each time point for treated and untreated samples. Bottom panel-Immunocytochemistry of LMB-treated and untreated Vero samples at 48 hours. Antibodies used were anti-NS5 (in green) and DAPI (in blue). Shown on the right is a graphical representation of the percentage of infected cells at each time point for treated and untreated samples. (B) Top panel-Immunocytochemistry of LMB-treated and untreated SK-N-SH samples at 48 hours. Antibodies used were anti-UPF1 (in green), anti-4G2 (in red), and DAPI (in blue). Shown on the right is a graphical representation of the percentage of infected cells at each time point for treated and untreated samples. Bottom panel-Immunocytochemistry of LMB-treated and untreated SK-N-SH samples at 48 hours. Antibodies used were anti-NS5 (in green) and DAPI (in blue). Shown on the right is a graphical representation of the percentage of infected cells at each time point for treated and untreated samples. Two images were counted for each antibody per sample and values were averaged. Error bars represent SEM.

The same experiment was carried out for ZIKV-infected SK-N-SH samples. At 4hpi for all antibodies, no positive signals were detected in any cell. By 24hpi, there were no major differences in positive signal for untreated (0.0%, 0.0%, 1.19%) versus treated (0.0%, 0.0%, 1.44%) cells for anti-4G2, -ZIKV NS5, and -ZIKV capsid, respectively. At 48hpi, there were notable differences in positive signal between untreated (19.51%, 16.45%, 26.64%) and treated (3.82%, 1.53%, 18.27%) cells for anti-4G2, -ZIKV NS5, and -ZIKV capsid, respectively (**Fig. 7B**). Additionally, treatment of 20nM LMB resulted in cytotoxicity for many treated SK-N-SH cells, an effect which has been reported to occur in neuroblastoma-derived cell lines upon LMB treatment (*45*). Taken together, while treatment with 20nM LMB exhibited changes in ZIKV infectivity over 48hpi, it did not completely inhibit ZIKV from infecting the Vero and SK-N-SH host cells. While nuclear export does play a role in ZIKV infectivity, host nuclear export is only one part of the overall viral barrage affecting the host cell.

## Discussion

In this study, we examined the dynamic changes occurring throughout the early, middle, and late stages of acute infections, as well as long term ZIKV and DENV infections. The key conclusions that can be drawn from our experiments are: 1. Transcriptomic and protein profiles distinguish exposure to ZIKV and DENV viruses in host cells. 2. Several host regulatory circuits are deregulated during ZIKV and DENV infections. Investigations revealed that most transcripts from the mRNA processing pathway are upregulated, with the dynamics of this up-regulation being cell type specific. Further analysis showed that transcripts related to NMD and RNA degradation complexes are upregulated. 3. The NMD pathway master regulator UPF1 exhibits increased transcript levels, however, at the protein level, they are degraded by ZIKV but not DENV infection. Another co-factor of NMD, UPF2, is also degraded upon ZIKV infection but not in DENV infection. 4. During acute infection with ZIKV, host cells accumulate intronic reads in many transcripts due to deregulation of the ‘NMD’ and ‘RNA degradation’ pathways. Intronic accumulation in some transcripts is also observed in DENV -infected cells but only after a long-term infection. 5. Interestingly, we found that NPC genes are also upregulated in a cell type specific manner upon ZIKV and DENV infection. 6. Inhibiting nuclear export with Leptomycin B prior to ZIKV exposure causes decreased ZIKV virus infection in host cells indicating the important role NPC complex plays in promoting flavivirus infection. Our data underscores the potential of targeting the NPC for developing novel anti-flavivirus therapeutics.

The virulence of a flaviviruses is the cumulative result of all virus-host interactions. Subtle genetic differences between flavivirus species can significantly alter the virulence and tissue tropism of the respective virus. As ZIKV and DENV invade host cells, a struggle for transcriptional control begins. Host signaling molecules begin alerting of invading pathogens and trigger up-regulation of genes associated with host defense. Despite the transcriptomic up-regulation of the host defense genes, ZIKV capsid-mediated degradation of the master regulators of the NMD pathway, UPF1 and UPF2, allows the virus to evade aspects of host defense and replicate more efficiently (**Fig. 8**) (*46, 47*). We believe that ZIKV-mediated degradation of UPF1 and UPF2 is one of the important factors that causes early cytopathy observed in Vero and SK-N-SH cells. Since DENV is unable to degrade these host proteins, it exhibits reduced virulence and little to no cytopathy at early stages of acute infection (*48*).

**Fig. 8.**
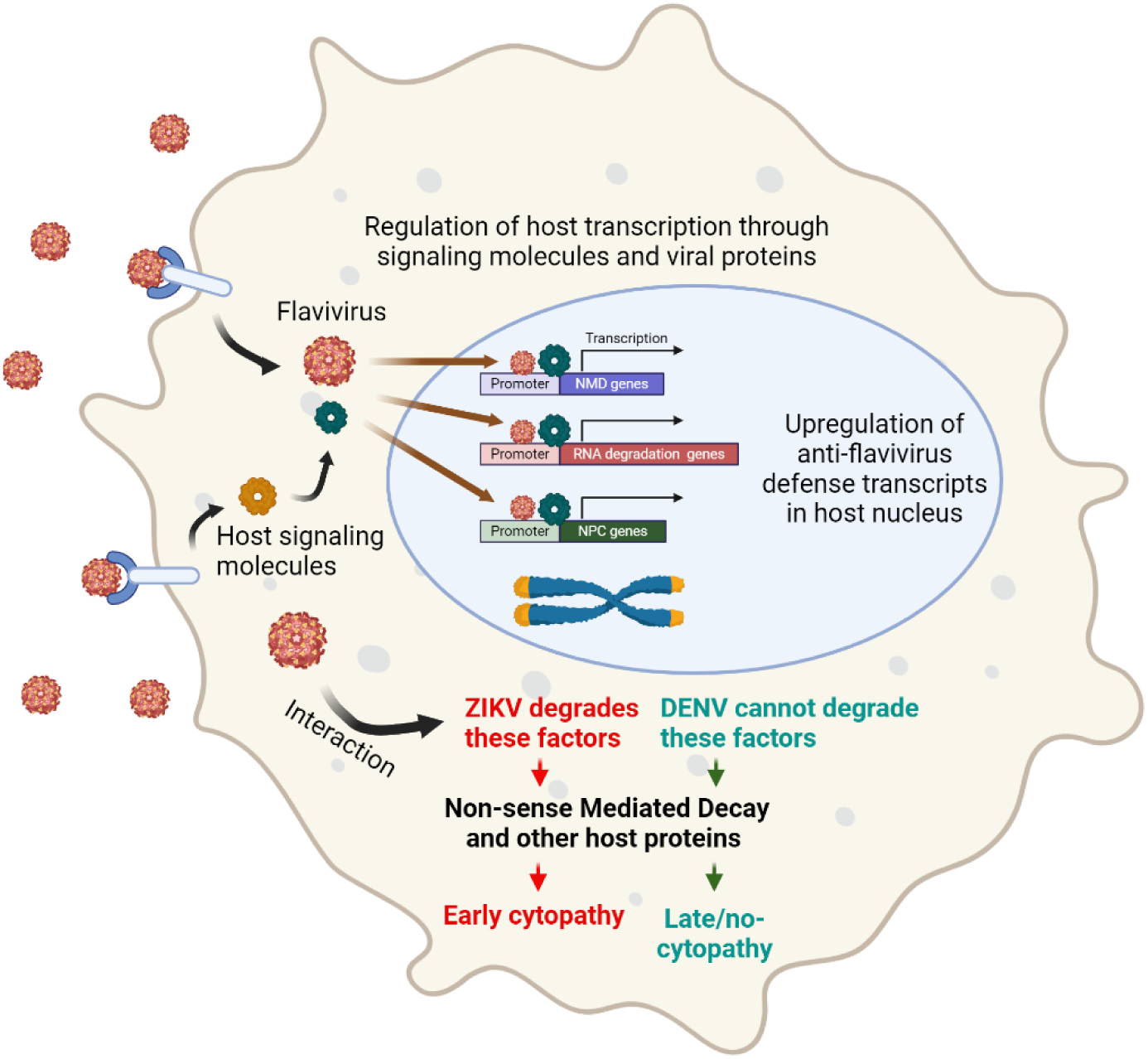
Proposed model of regulation of host defense with respect to ZIKV and DENV virulence. The diagram presents the interaction between ZIKV and DENV and host cells. As viruses invade host cells, host signaling molecules alert the cell to invading viral proteins which initiates altering host gene transcription and activating mRNA processing transcripts. ZIKV, but not DENV, viral proteins enter the cytoplasm where they degrade host nuclear UPF1 proteins to subvert host viral defenses, this host protein degradation further promotes ZIKV infection. As a result of degradation of UPF1 and other host proteins, nuclear mRNA transcripts begin to accumulate, and also leads to cytopathy in ZIKV-infected samples.

It has been reported that UPF1 associates with transcription sites and positively influences eukaryotic transcription (*49, 50*). Consistent with previous studies, the opposite trends observed in the transcriptional regulation of certain genes associated with the NMD complex, RNA degradation complex, and NPC complex on days 1 and 3 following ZIKV and DENV infection in Vero cells (downregulated in ZIKV cells and upregulated in DENV infected cells) might be due to the early reduction of UPF1 after ZIKV infection and the increase of UPF1 protein following DENV infection.

Due to the viral-induced protein degradation of master NMD regulators, there is an accumulation of intronic reads in ZIKV-, but not DENV-infected samples during acute infection. While the intronic accumulation of reads was most obvious in ZIKV-infected SK-N-SH samples, intronic reads accumulated in ZIKV-infected Vero samples as well (**Fig. S10C, 5C**). There were cases, such as with *NXT1*, where ZIKV-infected Vero exhibited intronic read accumulation, but no such accumulation was observed in ZIKV-infected SK-N-SH (**Fig. 6C, Fig. S11B)**. This difference in intronic read accumulation may be due to a difference in viral-host species interaction. No major differences were observed in intronic reads for long-term ZIKV infected Vero samples, suggesting that a balance may be restored over time. This is likely due to the accumulation of favorable mutations in ZIKV, allowing the host cell to partially recover UPF1 protein levels as the infection reaches its long-term phase.

Interestingly, we found that NPC transcripts followed a similar pattern to other ‘complex’ transcripts studied in this article. ZIKV infection did not reduce the NXT1 protein level. Further investigation using LMB identified the role of NPC in promoting flavivirus infection. It is well known that flavivirus proteins, such as NS5, shuttle to the eukaryotic nucleus, which is required for viral replication (*51*). Our study and a previous study indicates that ZIKV has developed the unique ability to degrade host NMD factors (*52*) to protect itself from host defenses and leverage the NPC to swiftly shuttle to the eukaryotic nucleus and promote its replication. Finally, this study and other previous studies (*28–30, 36, 52*) have uncovered several complexes, such as NMD complex, that play a pivotal role in the defense against flavivirus infections in host cells. It would be intriguing to explore whether transcripts linked to these complexes, or their target transcripts, could serve as biomarkers for detecting flavivirus infections in cells and tissues intended for therapeutic product production.

### Limitation of the study

In this article, we demonstrate that ZIKV and DENV induce cell-type-specific up-regulation of key transcripts linked to certain regulatory complexes, while ZIKV uniquely causes degradation of specific host proteins. However, our findings, derived from two cell lines, may not comprehensively reflect the transcriptional and protein changes these viruses induce at the organismal level. Understanding how host defense mechanisms interact across various cell types in response to ZIKV and DENV virulence is crucial for a more complete picture. Additionally, protein changes were assessed using western blot and ICC assays, which do not reflect the complete proteome dynamics during ZIKV and DENV infection. Furthermore, our LMB study focused solely on ZIKV infection in these cell lines, necessitating a more comprehensive investigation to evaluate the potential of targeting NPC complexes for antiviral therapy against flaviviruses.

## Materials and Methods

### Experimental Design

This study aimed to uncover host molecular factors that regulate the virulence of ZIKV and DENV. Acute and persistent flavivirus-infected cultures were prepared, and samples were collected at various time points. To determine transcriptomic and protein changes, RNA-seq and protein-based assays were performed on mock- and flavivirus-infected human and green monkey samples. RNA-seq analysis identified cell-type and time-dependent upregulation of mRNA processing factors such as NMD, RNA degradation, transcripts as well as NPC transcripts. Nuclear export was inhibited by 20nM leptomycin B to determine whether active nuclear transport influenced ZIKV virulence.

### Cell culture

Cell lines used include Vero E6, and SK-N-SH. Cultures of Vero and SK-N-SH, were performed as previously described (*53*). Cultures were grown in an incubator at 37°C in 5% CO_2_. Cell cultures were grown in triplicate wells for each experiment. Cultures were grown in RPMI 1640 (Vero) or MEM (SK-N-SH) media supplemented with 10% fetal bovine serum (FBS) (Cat# A15-201, PAA Laboratories) and Penicillin-Streptomycin (P/S) (Cat# 30-002-CI, Corning) in 6-well plates until 80-90% confluent. Cell % viability was determined using a Countess 3 automated cell counter with 0.4% Trypan Blue solution (Cat# 15250061, Gibco). At least 2 counts per sample were taken and averaged.

### Generation of acute and long-term infected cell lines

Three wells of each culture plate with cells at 80-90% confluence was acutely infected with ZIKV MR766 or DENV-3 at Multiplicity of Infection (MOI) of 1 in respective media supplemented with 2% FBS + P/S for 2 hours. The remaining three wells of each culture plate were treated with comparable volume of supernatant from healthy Vero or SK-N-SH cells (Mock/control) in respective media supplemented with 2% FBS + P/S. Following acute infection (6+ DPI), cells were grown in media supplemented with 10% FBS + P/S and passaged weekly for at least 3 months while still exhibiting positive signal for anti-flavivirus antibodies in ICC assay to be considered as long-term infected (**Fig. S1**). During acute infection, samples were collected, and cells were counted using a Countess 3 (Thermo Fisher) daily for 5 days. Samples were prepared for RNA and protein analysis prior to storing in −80°C. Slides for immunocytochemistry were prepared and fixed in acetone:methanol (1:1) for 10 minutes at room temperature.

### Western blot and immunocytochemistry assays

Pellets from mock, acute, and long-term ZIKV MR766 and DENV-3 infections were resuspended in RIPA buffer (Cat# 89901, Thermo Scientific) containing cOmplete Mini protease inhibitor cocktail (Cat# 11 836 153 001, Roche) for 1 hour on ice. Samples were sonicated (sonicator Model Q500, QSONICA) with a cup horn attachment (Cat# 431C2, QSONICA) at 40% power for 3 minutes (30 seconds on/off). Protein concentrations were determined via bicinchoninic acid (BCA) assay (Cat# 23225, Thermo Fisher) performed in duplicate. After normalizing the sample concentrations, the samples were treated with 5% 2-mercaptoethanol in Laemmli buffer and heated at 95°C for 5 minutes prior to loading into a 4-15% gradient SDS gel (Cat# 4568085, Bio-Rad) and run at 160v for 50-55 minutes. The gels were transferred to PVDF membranes (Cat# IB24002, Invitrogen) using an iBlot2 system set to 25v for 7 minutes. The transferred membranes were cut, washed, and blocked in 5% nonfat milk that was dissolved in 0.1% tween20 (Cat# 170-6531, Bio-Rad) in TBS (Cat# 1706435, Thermo Scientific) (TBS-T). Membranes were rinsed and incubated in primary antibody for 1 hour at room temperature (RT), following which, they were washed 3x for 5 minutes each in TBS-T. Membranes were incubated in secondary antibody for 30 minutes at RT and then washed 3x for 5 minutes each in TBS-T. The membranes were developed in Pico western blot developing reagent (Cat # 34580, Thermo Scientific) or Femto western blot developing reagent (Cat #34096, Thermo Scientific) and imaged on an Amersham Imager 600. Membranes were stripped using Restore PLUS Western Stripping Buffer (Cat# 46430, Thermo Scientific), washed with TBS-T, blocked, and subjected to a new antibody probe as previously described.

Various antibodies were used in western blot and ICC. Primary antibodies used were as follows: Mouse (M) anti-4G2 (Cat# Ab00230-2.0, Absolute Antibody), M anti-DENV NS1 (Type 2/3/4) (Cat# MAB94441-100, R&D Systems), M anti-NXT1 (Cat#67680-1-Ig, Proteintech), M anti-alpha tubulin (Cat#66031-1-Ig, Proteintech), Rabbit (Rb) anti-beta actin (Cat# 026-42210, Li-cor), Rb anti-DENV NS5 (Cat#NBP2-42901, Novus Biologicals), Rb anti-GAPDH (Cat# 5174S, Cell Signaling Technology (CST)), Rb anti-UPF1 (Cat#12040, CST) Rb anti-UPF2 (Cat#11875, CST), Rb anti-ZIKV NS5 (Cat# GTX133312, GeneTex), Rb anti-ZIKV envelope (Cat# GTX133314, GeneTex). Secondary antibodies used were as follows: Peroxidase-labeled goat anti-rabbit IgG (H+L) (Cat# 5450-0010, KPL), peroxidase-labeled goat anti-mouse IgG (H+L) (Cat# 214-1806, KPL), goat anti-mouse AlexaFluor 488 Plus (Cat# A32723, Invitrogen), goat anti-mouse AlexaFluor 568 Plus (Cat# A11004, Invitrogen), goat anti-rabbit AlexaFluor 488 Plus (Cat# A32731, Invitrogen), and goat anti-rabbit AlexaFluor 568 Plus (Cat# A11011).

### Preparation of Total RNA, rRNA-depletion, and generation of cDNA libraries

Total RNA was extracted from samples using a combination of TRIzol extraction and spin column-based purification. RNA from mock, acute, and long-term infections were treated with 500µl TRIzol reagent (Cat# 2050-1-200, Zymo Research). Samples were incubated for 5 minutes on ice and 100µl chloroform (Cat# J67241.K2, Thermo Scientific) was added. Samples were mixed by inverting and incubated for 3 minutes at RT, then centrifuged for 15 minutes at 12,000 RPM at 4°C. The clear supernatant was collected in a new tube and an equal amount of 100% isopropanol was added. The samples were mixed by inverting and incubated for 10 minutes at 4°C to precipitate the total RNA. Samples were transferred into a PureLink RNA Mini (Cat# 12183025, Invitrogen) column and RNA was purified according to manufacturer’s instruction. Total RNA concentration was determined via a Qubit 4 fluorometer with a Qubit RNA High Sensitivity kit (Cat# Q32852, Invitrogen), and RNA quality was checked using Agilent 2100 Bioanalyzer with an RNA 6000 nano kit (Cat# 5067-1511, Agilent).

A NEBNext rRNA Depletion Kit (Human/Mouse/Rat) (Cat# E6310X, New England Biolabs) was used to deplete rRNA from the previously prepared total RNA. Samples were rRNA-depleted as described in Section 1 of the respective kit manual. RNAClean XP beads (Cat# A63987, Beckman Coulter) were used for recovery of RNA during rRNA depletion. Preparation of cDNA libraries was performed using NEBNext Ultra II RNA Library Kit for Illumina (Cat# E7770L, New England Biolabs) following Section 4 of the respective manual. AMPure XP beads (Cat# A63881, Beckman Coulter) were used for cDNA recovery of prepared libraries. NEBNext Multiplex Oligos for Illumina (Index Primer Sets 1-4) (Cat# E7335L, E7500L, E7710L, E7730L, New England Biolabs) were used for indexing during library preparation. Prepared libraries were checked for quality using Agilent 2100 Bioanalyzer prior to sequencing. Sequencing was performed at the FDA/CBER Facility for Biotechnology Resources (FBR) Core on a NovaSeq 6000 using a S1 flow cell.

### TCID_50_

Infectious virus titers were determined by performing TCID50 assays on Vero-E6 cells (*54, 55*). TCID_50_ was performed by seeding 1 x 10^4^ Vero cells in each well of a 96-well plate and growing to 80-90% confluence with ∼100% viability in RPMI1640 supplemented with 10% FBS + P/S. Serial dilutions of mock, acute, and long-term ZIKV and DENV viral supernatants were prepared in RPMI1640 supplemented with 2% FBS + P/S. Media was removed from 96-well plates and viral serial dilutions (8 replicates per dilution) in media were added and incubated at 37°C for 5 days. After 5 days, wells were checked for 50% CPE and results were tallied for each dilution and infection type. TCID_50_ values were calculated for each infection type using the Spearman–Kärber algorithm (*56*).

### Inhibition of nuclear export during ZIKV infection via Leptomycin B

Vero and SK-N-SH cells were grown to 80 to 90% confluence in 12 well plates. Leptomycin B (20nM) was added to RPMI1640 supplemented with 2% FBS + P/S. Respective cell cultures were incubated in LMB-supplemented media for 3 hours prior to introduction of ZIKV MR766 at MOI=1. Cell pellets were collected at 4-, 24-, and 48-hpi, and slides were fixed in acetone:methanol (1:1) and prepared for ICC assays.

### Quality assessment and read filtering

FastQC (https://www.bioinformatics.babraham.ac.uk/projects/fastqc/) and MultiQC (*57*) was used to assess the quality of the raw FASTQ data using its default options. The assessment revealed high data quality. Notably, adapter sequences were detected within the dataset, prompting the need for preprocessing steps. To prepare the data for downstream analysis, fastp (*58*) was employed to trim adapter sequences, trim polyG in read tails, and filter out low-quality reads. Reads with a Phred score below 15 were discarded, along with those shorter than 5 bases.

### Genome mapping using HISAT2 and STAR

Alignment of the preprocessed reads to the *C. sabaeus* reference genome (chlSab2) and human genome was conducted utilizing both HISAT2 (*59*) and STAR (*60*) alignment tools. The genome FASTA file of chlSab2 was downloaded from UCSC website (https://hgdownload.soe.ucsc.edu/goldenPath/chlSab2/bigZips/). The genome indices were built using “*hisat2-build*” and “*STAR –runMode genomeGenerate*” commands with the default options.

### Quantification of reads per gene

The quantification of human and *C. sabaeus* reads per gene was performed using featureCounts (*61*) for both HISAT2 and STAR mapping results. Additionally, STAR’s GeneCounts function was employed specifically for the STAR mapping results, resulting in the generation of three distinct read-count datasets. Considering the impact of mapping and quantification methods on the count numbers, particularly for genes with low expression levels, comparative analysis of the three read-count datasets was carried out to identify genes exhibiting big read count variation exceeding two folds between the three datasets.

For *C. sabaeus* data, approximately 34% of the mapped reads remained unassigned to annotated gene regions. Further analysis revealed that many of these reads mapped to non-annotated regions, where homologous genes are present in other primate genomes.

### Screening for viral origins and green monkey sequences

Unmapped reads were investigated for viral origins of ZIKV MR766 and DENV3, through alignment using BWA (*62*). The FASTA file of ZIKV MR766 was downloaded from UCSC website (https://hgdownload.soe.ucsc.edu/hubs/GCF/000/882/815/GCF_000882815.3/), and the FASTA file of DENV3 was downloaded from UCSC website (https://hgdownload.soe.ucsc.edu/hubs/GCF/000/866/625/GCF_000866625.1/). Reads unaligned to the latest version of green monkey genome, ZIKV MR766, or DENV3 underwent de novo assembly using SPAdes (*63*). Subsequent BLAST (https://blast.ncbi.nlm.nih.gov/Blast.cgi?PROGRAM=blastn&PAGE_TYPE=BlastSearch&LINK_LOC=blasthome) analysis of assembled contigs revealed imperfect matches to primate genomes, including human, rhesus monkey, gorilla, and chimpanzee.

### Secondary analysis of *C. sabaeus* data

First, reads quality was assessed using fastp version 0.23.4 to trim adapter sequences. Reads were aligned to the *C. sabaeus* reference genome Vero_WHO_p1.0 and ZIKV reference using HISAT2 version 2.2.1 and STAR version 2.7.11b, which generated binary alignment map (BAM) files. Read metrics were collected using Picard version 3.2.0. featureCounts version 1.5.3 was used to report the count of alignments from each sample’s BAMs at the universe of regions and generate a counts matrix which was used in downstream analysis.

### Differential expression analysis and pathway analysis

R/DESeq2 (*64*) were used for differential expression analysis by comparing virus-treated human and *C. sabaeus* samples with the according mock samples across different treatment durations (1-, 3-, and 5-DPI). Additionally, the Vero long term infected samples were compared with all mock samples for the differential expression analysis. The alpha and the fold change thresholds were set to a Benjamin–Hochberg-adjusted p-value <0.05 and 1.5 respectively for significance in the DGE test. Read counts were normalized using the median of ratios method. DGE cluster analysis was performed using the pheatmap R package.

The AnnotationForge, ensembldb, and biomaRt R packages were used to create a custom annotation database for *C. sabaeus*. The clusterProfiler package in R (*65*) was used to identify the Gene Ontology (GO) pathways (*66, 67*) enriched in the data, as well as to explore their potential biological processes, cellular components, molecular functions, and important signaling pathways. A minimum gene set of 10 and a maximum gene set of 500 were chosen. P < 0.05 and false detection rate < 0.05 were considered statistically significant for Gene Set Enrichment Analysis (GSEA) (*68*).

## Acknowledgments

The authors thank Drs. Fei Mo, Nirjal Bhattarai, and Carolyn Laurencot for critical reading of this manuscript and helpful comments. The authors thank Dr. Charles Chung for his help in creating custom annotation database for *C. sabeus* gene ontology enrichment analysis.

## Author Contributions

Conceptualization: AS, SD

Methodology: AS, BG, BL, NT, SK, JX, SD

Investigation: AS, BG, TF, BL

Visualization: AS, SD

Supervision: JX, SD

Writing- original draft: AS, BG, NT, JX, SD, SK

Writing- review & editing: AS, BG, MR, JX, SD, SK

## Data availability statement

All data needed to evaluate the conclusions in the paper are present in the paper and/or the Supplementary Materials.

The ZIKV and DENV long-term infected Vero cells can be provided by CBER/FDA pending scientific review and a completed material transfer agreement. Requests for the long-term infected cells should be submitted to: Sandip De.

## Supplementary information

**Table S1**. GSEA adjusted p-values for mRNA processing (GO:0006397), mRNA metabolic process (GO:0016071), and RNA splicing (GO:0008380) pathways in mock versus ZIKV-, and DENV-infected Vero samples as well as mock versus ZIKV-infected SK-N-SH samples.

**Table S2**. Immunocytochemistry assay percent infection cell counts for 20nM LMB-treated and untreated Vero and SK-N-SH infected with ZIKV (MOI=1).

**Fig. S1.**
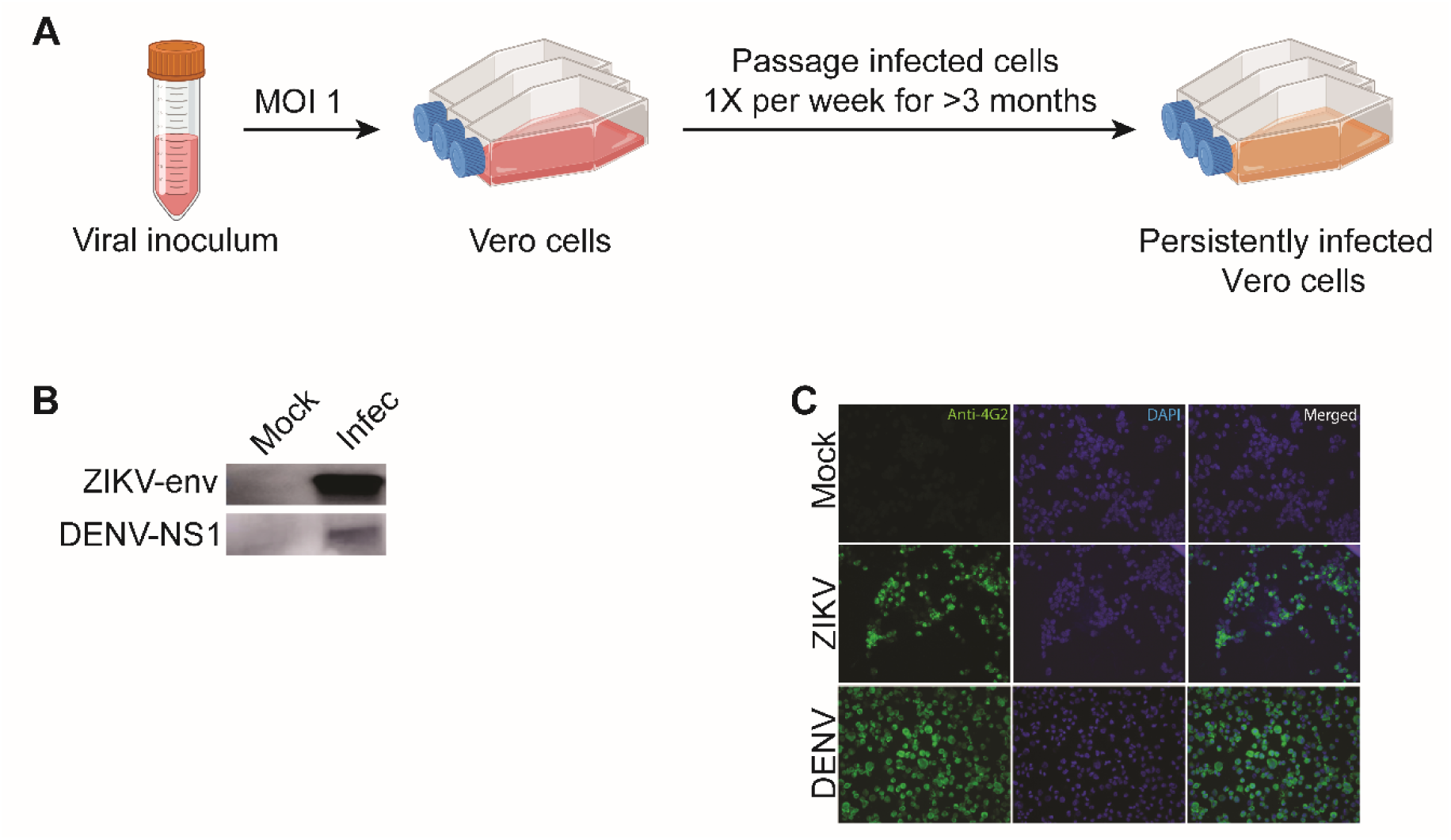
Generation of long-term infected cell lines. (A) Viral inoculums were introduced into Vero cell lines at MOI of 1. These cultures were passaged once per week for more than 3 months before testing for continued flavivirus presence. (B) Western blot analysis of mock, ZIKV-, and DENV-long-term infected Vero samples. Antibodies used were anti-ZIKV envelope and anti-DENV NS1 for the respective long-term infected lines. (C) Immunocytochemistry of Vero control and long-term infected ZIKV, and DENV samples. Antibodies used were anti-4G2 (anti-flavivirus, in green) and DAPI (in blue).

**Fig. S2.**
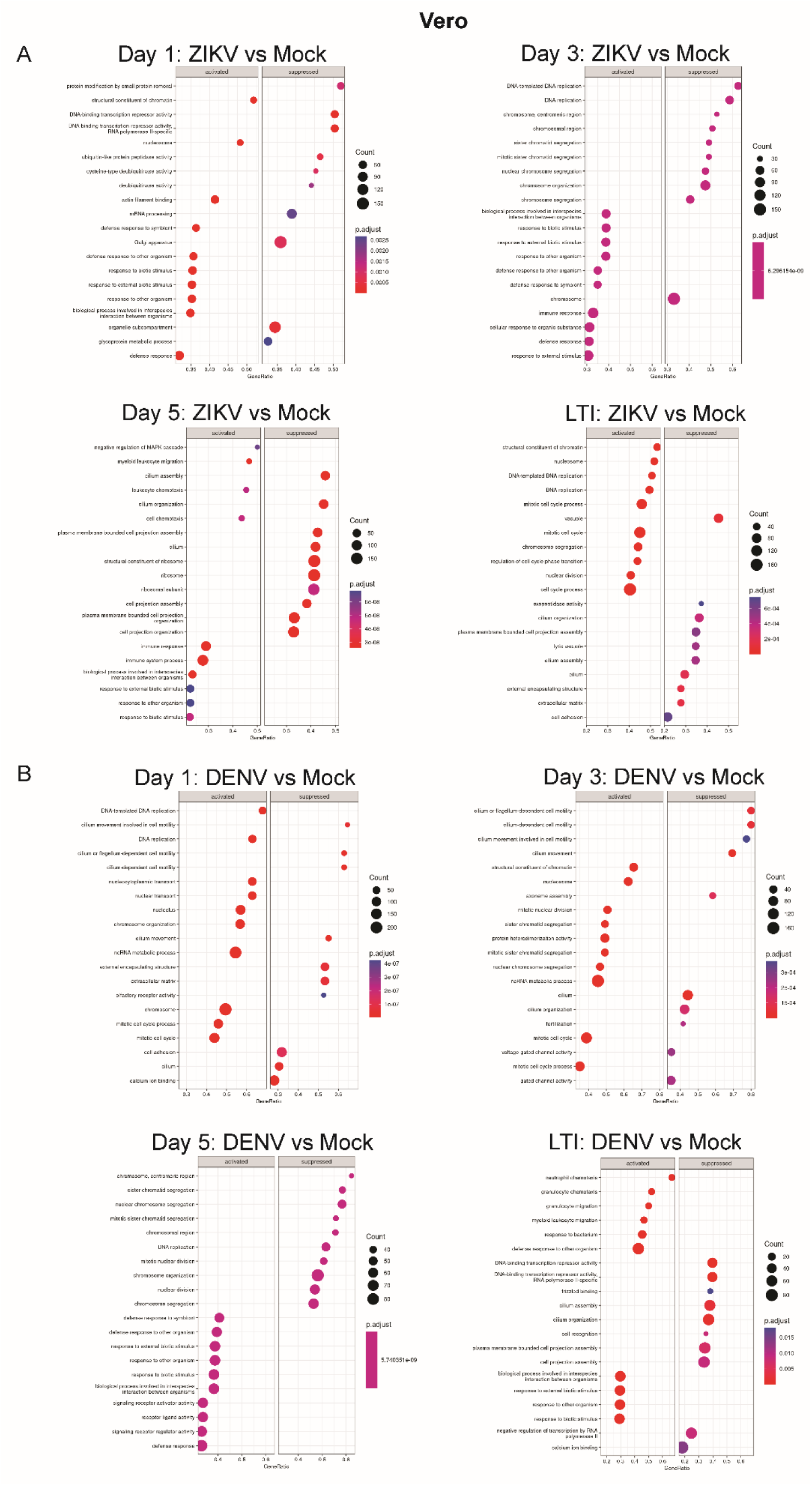
Vero mock versus acute and long-term ZIKV and DENV infection pathway analyses via GSEA. (A) A comparison of Vero acute and long-term ZIKV GSEA data, sorted to include only the top activated and suppressed pathways. (B) A comparison of Vero acute and long-term DENV GSEA data, sorted to include only the top activated and suppressed pathways.

**Fig. S3.**
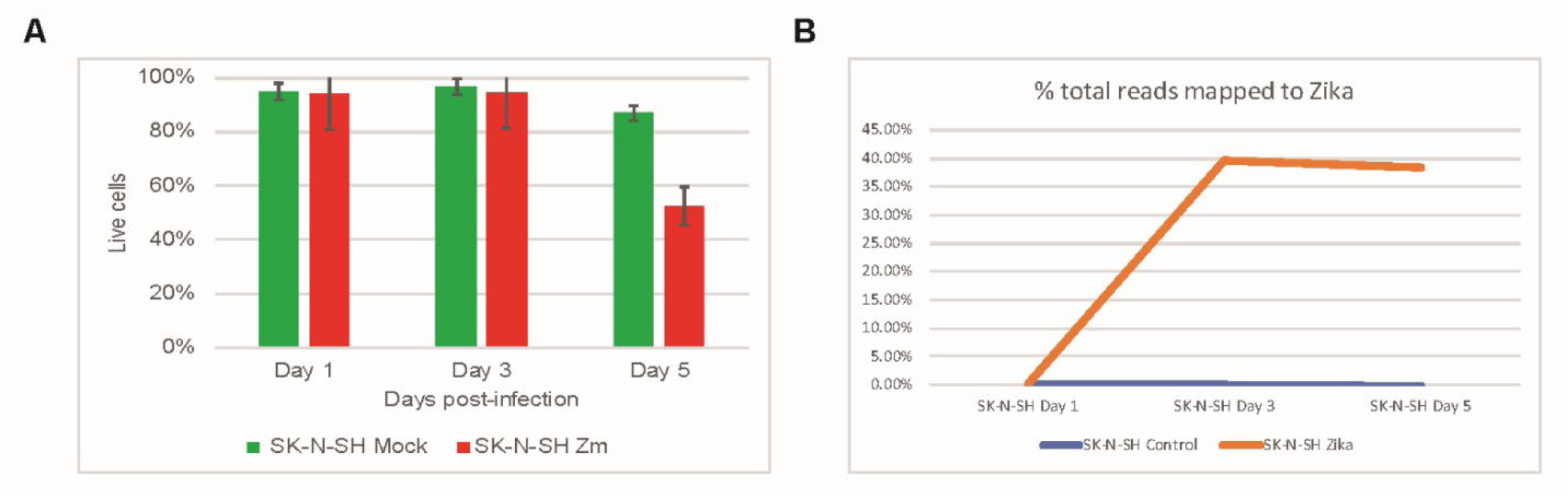
ZIKV-infected SK-N-SH cell viability and RNA-seq reads mapped to ZIKV share similar trends with ZIKV-infected Vero samples. (A) Percent cell viability for mock-infected control (green) and SK-N-SH ZIKV (red) samples on 1-, 3-, and 5-DPI. (B) Percent of total RNA-seq reads that mapped to ZIKV-MR766 (orange-infected SK-N-SH and blue-mock SK-N-SH samples) at 1-, 3-, and 5-DPI.

**Fig. S4.**
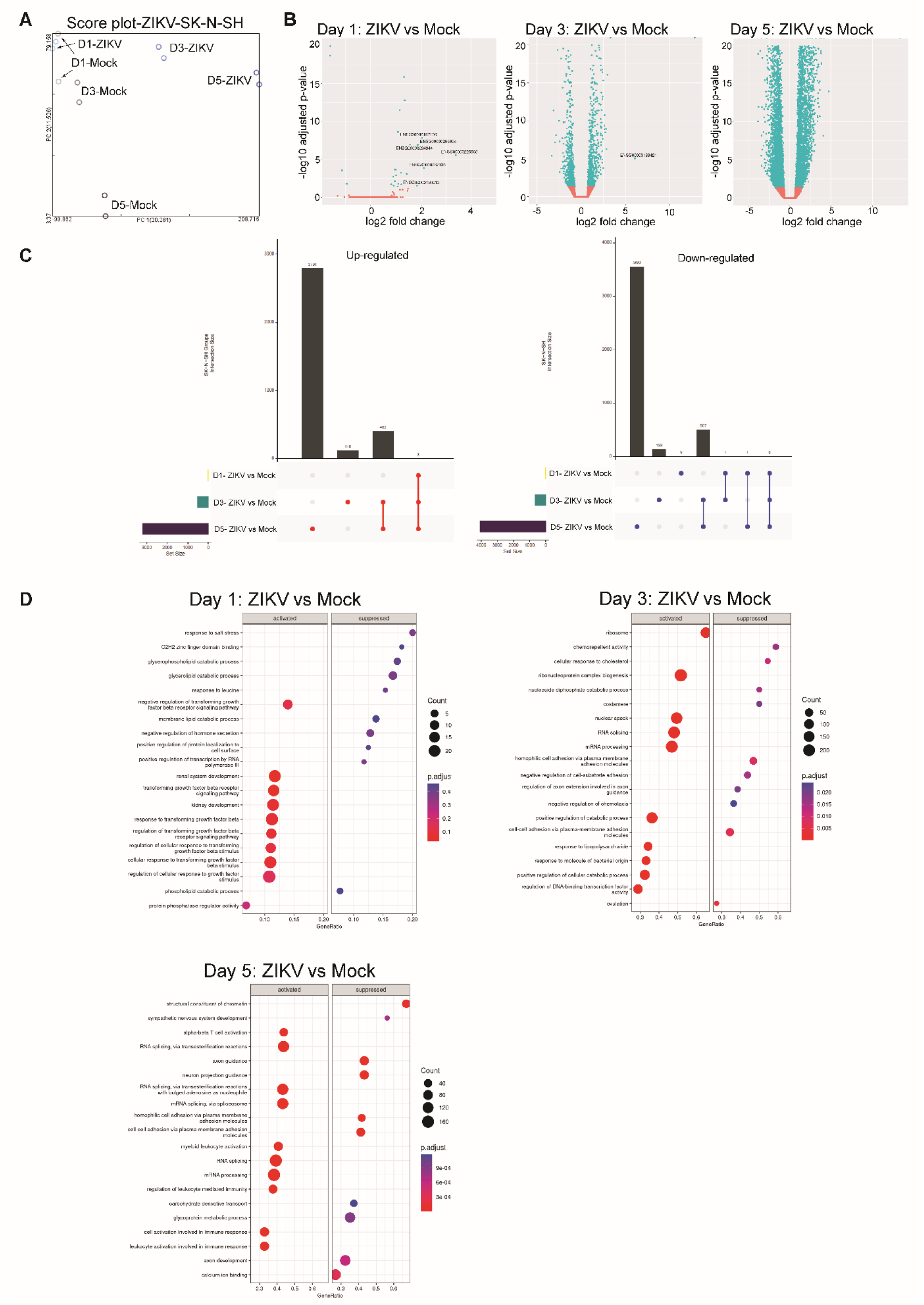
SK-N-SH mock versus acute ZIKV infection volcano plot and pathway analyses via GSEA. (A) PCA of mock, and ZIKV infected Vero samples for 1-, 3-, and 5-DPI used in RNA-seq analyses. Two replicates are graphed per condition per time point. (B) Volcano plots of −log_10_ adjusted p-value versus log_2_ fold change of differentially expressed genes between mock versus ZIKV-infected samples at 1-, 3-, and 5-DPI. DEGs with adjusted p-value >0.05 are shown in red while DEGs with adjusted p-value <0.05 are shown in teal. (C) UpSet plots of up-regulated and down-regulated genes shared between ZIKV infected vs mock Vero sample datasets from different days. (D) A comparison of SK-N-SH acute ZIKV infection GSEA data, sorted to include only the top 20 activated and suppressed pathways.

**Fig. S5.**
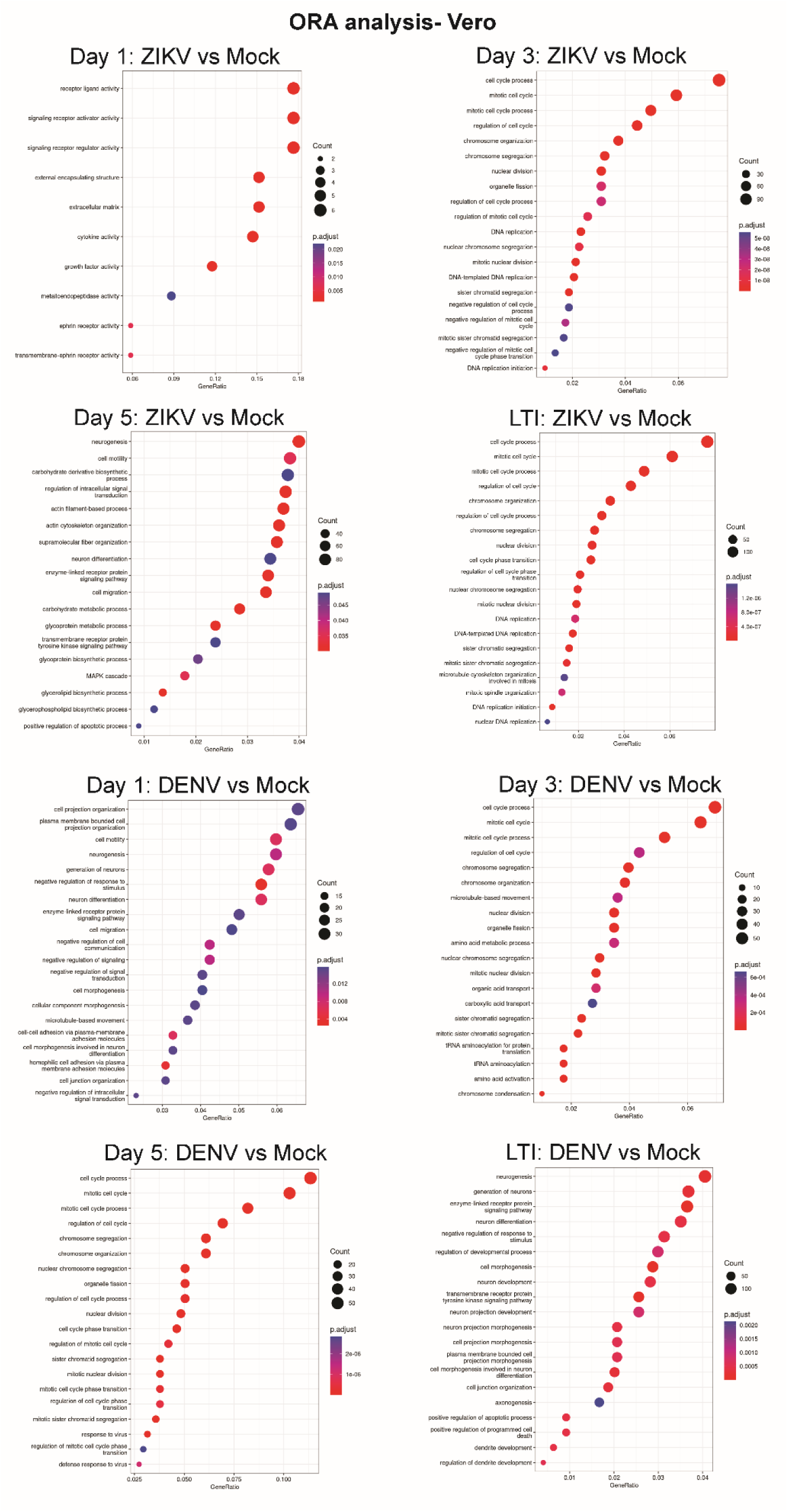
Vero mock versus acute and long-term ZIKV and DENV infection pathway analyses via ORA. A comparison of Vero acute and long-term ZIKV and DENV ORA data, sorted to include only the top differentially expressed pathways.

**Fig. S6.**
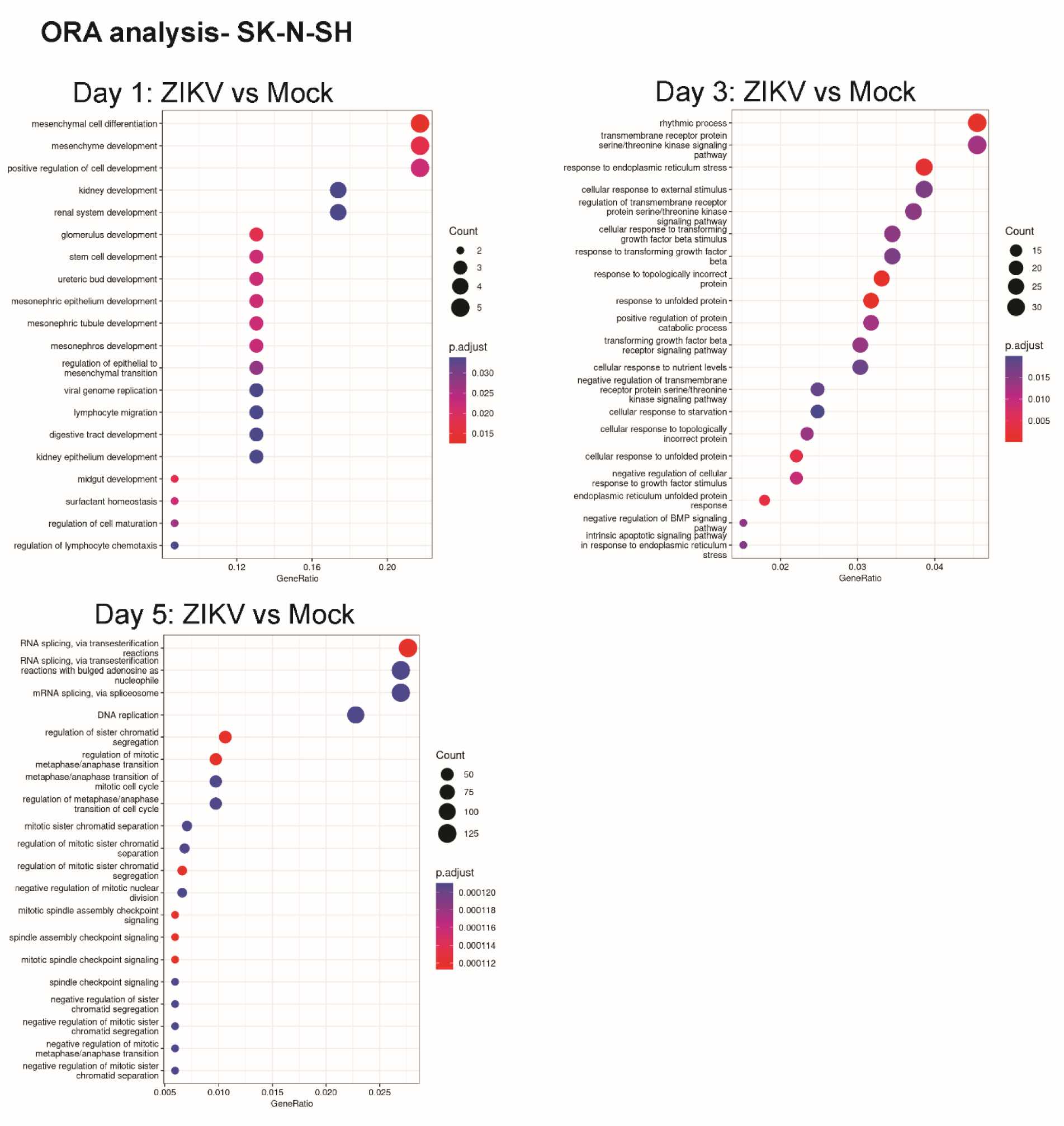
SK-N-SH mock versus acute ZIKV infection pathway analysis via ORA. A comparison of SK-N-SH acute ZIKV infection vs mock ORA data on different days, sorted to include the top 20 differentially expressed pathways.

**Fig. S7.**
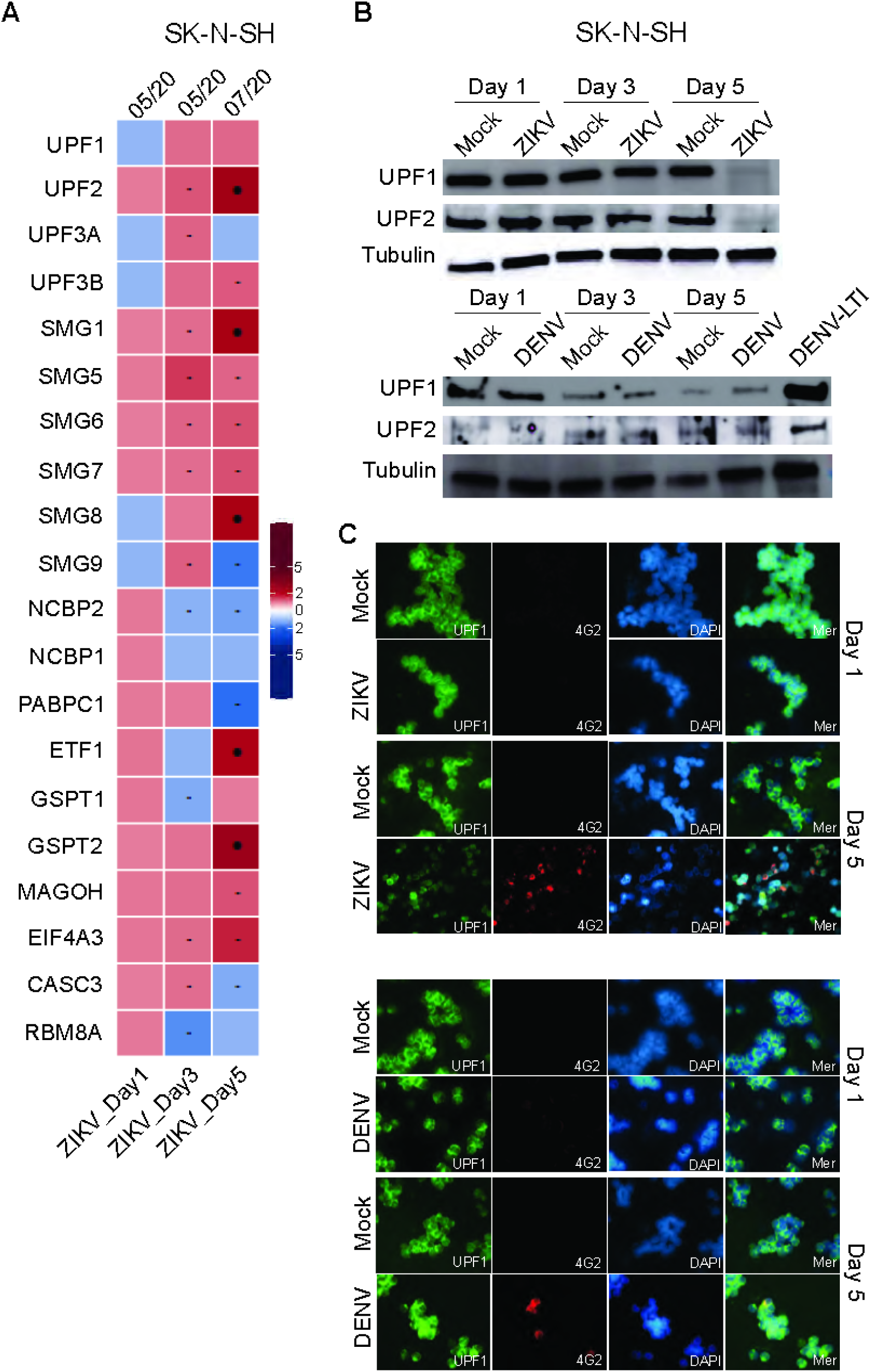
Investigation of NMD pathway transcripts in SK-N-SH ZIKV infections. (A) Heatmap of normalized RNA-seq data for mock versus ZIKV-infected SK-N-SH samples at 1-, 3-, and 5-DPI. (B) Western blot analysis of anti-UPF1, anti-UPF2, and anti-alpha tubulin for mock, ZIKV-, and DENV-infected SK-N-SH at 1-, 3-, and 5-DPI and long-term DENV infections. (C) Immunocytochemistry staining of SK-N-SH mock, ZIKV, and DENV at 1- and 5-DPI. Antibodies used were anti-UPF1 (in green), anti-4G2 (anti-flavivirus, in red), and DAPI (in blue).

**Fig. S8.**
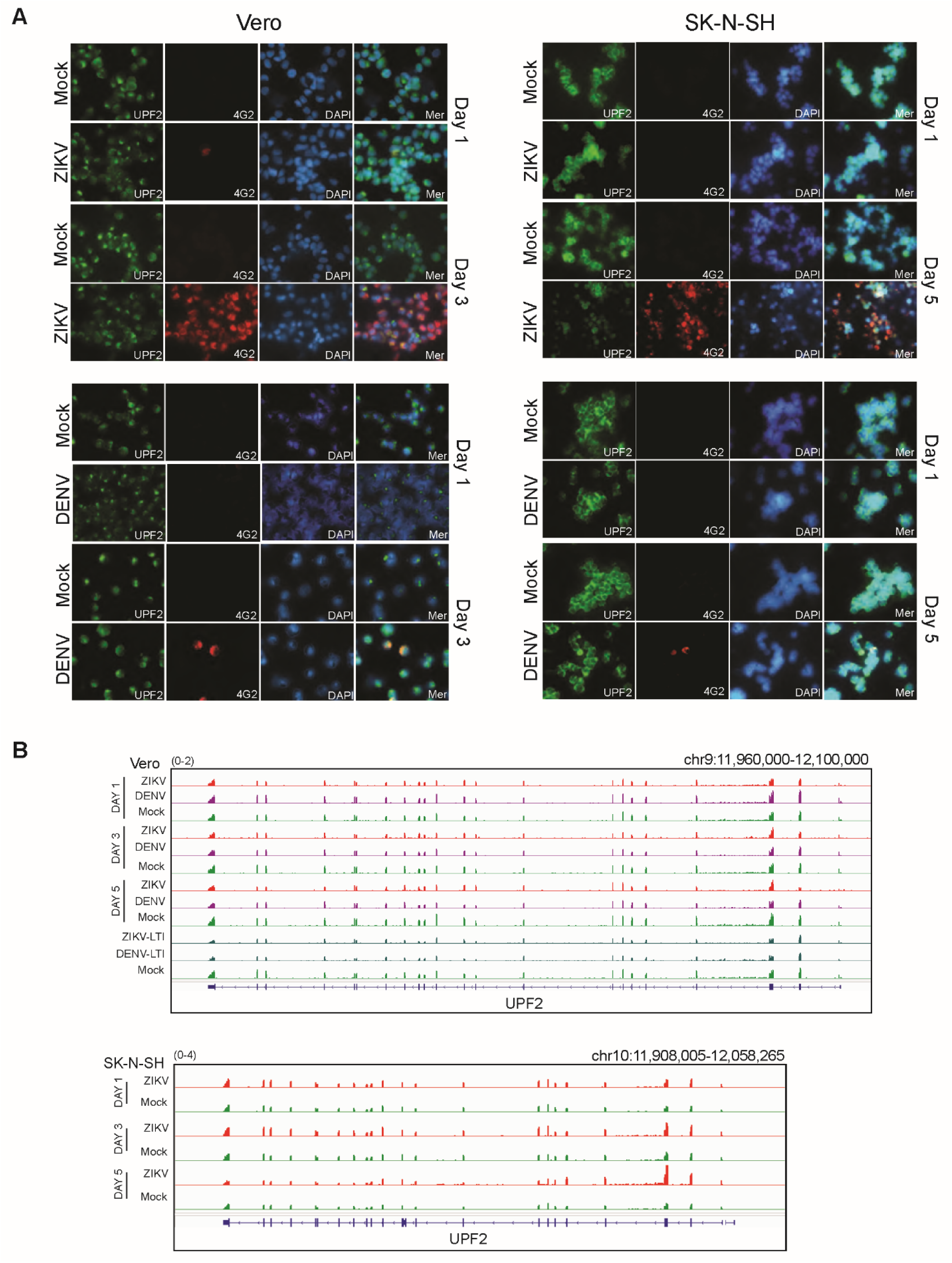
Immunocytochemistry of Vero and SK-N-SH mock versus acute ZIKV and DENV infection for UPF2. (A) Top panel-Immunocytochemistry of Vero and SK-N-SH mock, ZIKV-infected acute samples at days 1, 3 for Vero and days 1, 5 for SK-N-SH. Antibodies used were anti-UPF2 (in green), anti-4G2 (anti-flavivirus, in red), and DAPI (in blue). Bottom panel-Immunocytochemistry of Vero and SK-N-SH mock, DENV-infected acute samples at days 1, 3 for Vero and days 1, 5 for SK-N-SH. Antibodies used were anti-UPF2 (in green), anti-4G2 (anti-flavivirus, in red), and DAPI (in blue). (B) A comparison of normalized RNA-seq reads represented in reads per million for UPF2 and visualized in IGV. Comparisons shown for mock, ZIKV-, and DENV-infected Vero and SK-N-SH samples at 1-, 3-, 5-DPI, and long-term infected samples.

**Fig. S9.**
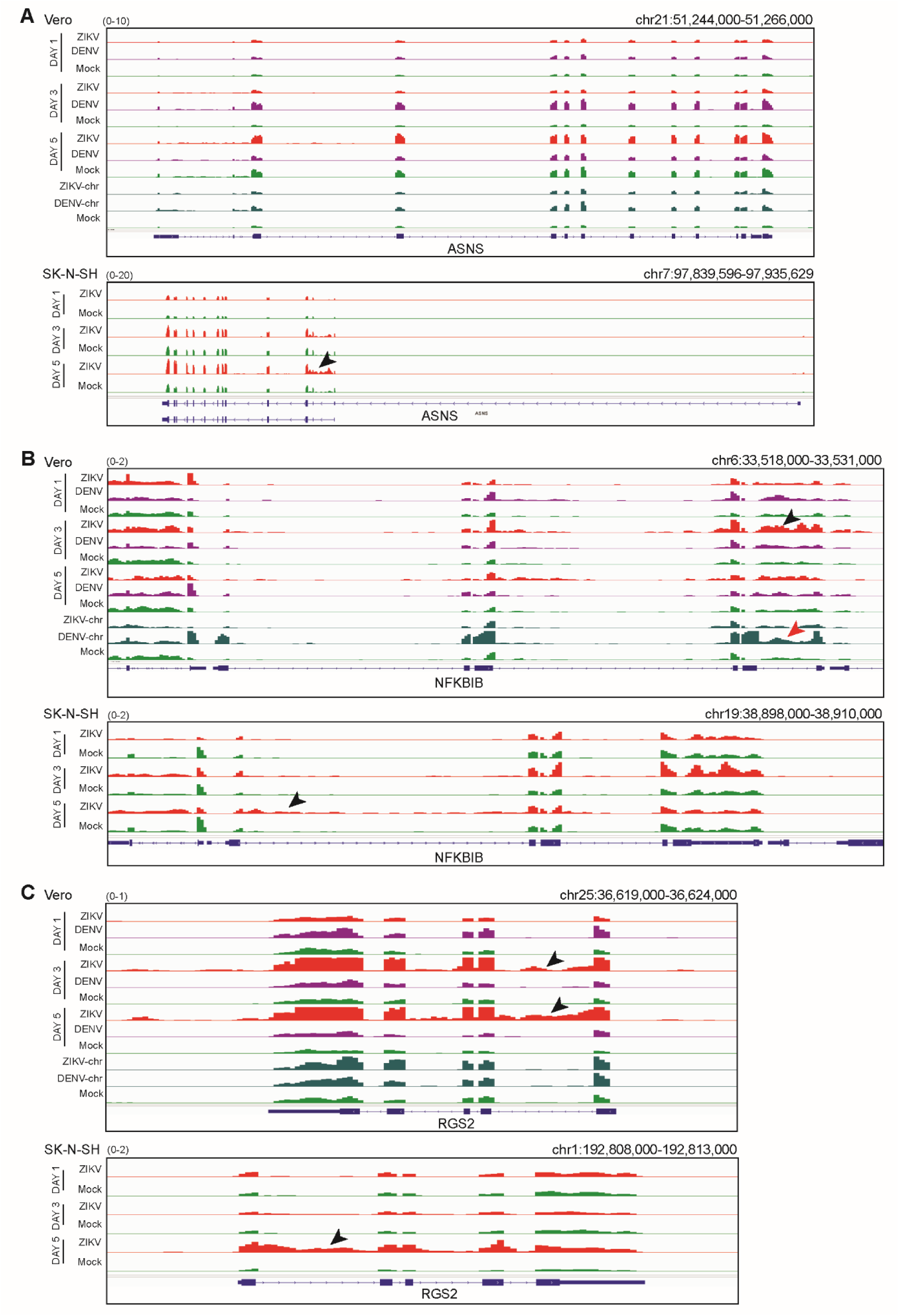
Comparison of Vero and SK-N-SH flavivirus infection data shows accumulated intronic reads during ZIKV infection. (A) A comparison of normalized RNA-seq reads represented in reads per million for *ASNS* and visualized in IGV. Comparisons shown for mock, ZIKV-, and DENV-infected Vero and SK-N-SH samples at 1-, 3-, 5-DPI, and long-term infected samples. Black arrowhead indicates notable accumulation of intronic reads. (B) A comparison of normalized RNA-seq reads represented in reads per million for *NFKBIB* and visualized in IGV. Black (ZIKV) and red (DENV) arrowheads indicate notable accumulation of intronic reads. (C) A comparison of normalized RNA-seq reads represented in reads per million for RGS2 and visualized in IGV. Black arrowheads indicate notable accumulation of intronic reads.

**Fig. S10.**
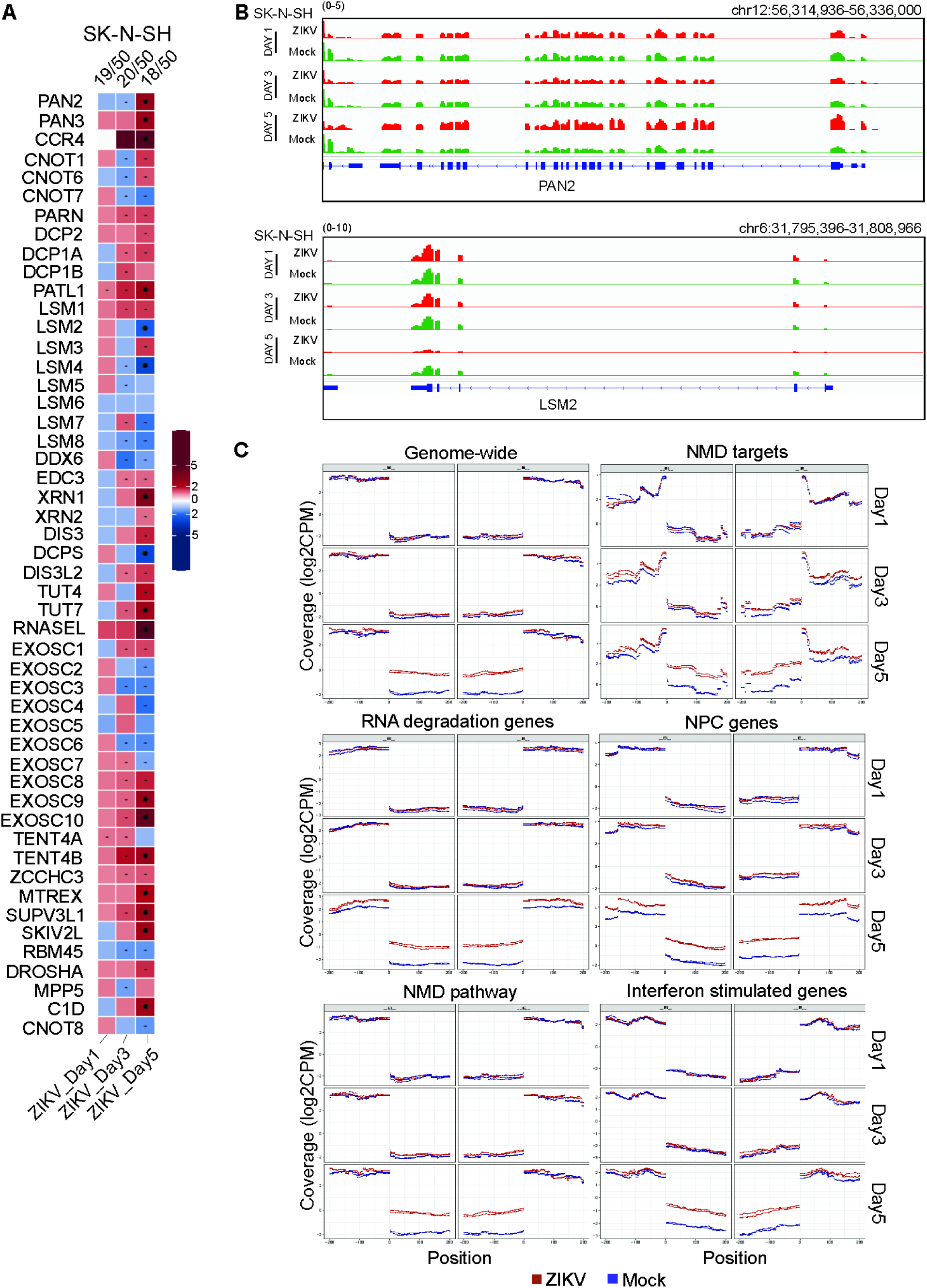
Accumulation of intronic reads observed in SK-N-SH cells during ZIKV infection. (A) Heatmap of RNA degradation-associated normalized RNA-seq transcripts for mock versus ZIKV-infected SK-N-SH samples at 1-, 3-, and 5-DPI. (B) A comparison of normalized RNA-seq reads represented in reads per million for PAN2 and LSM2 and visualized in IGV. Comparisons shown for mock and ZIKV-infected SK-N-SH samples at 1-, 3-, 5-DPI. (C) Comparison of intron/exon and exon/intron boundaries genome-wide as well as for NMD pathway, RNA degradation genes, NPC genes, NMD targets, and ISGs within 200bp for mock (blue) and ZIKV-infected (red) SK-N-SH samples.

**Fig. S11.**
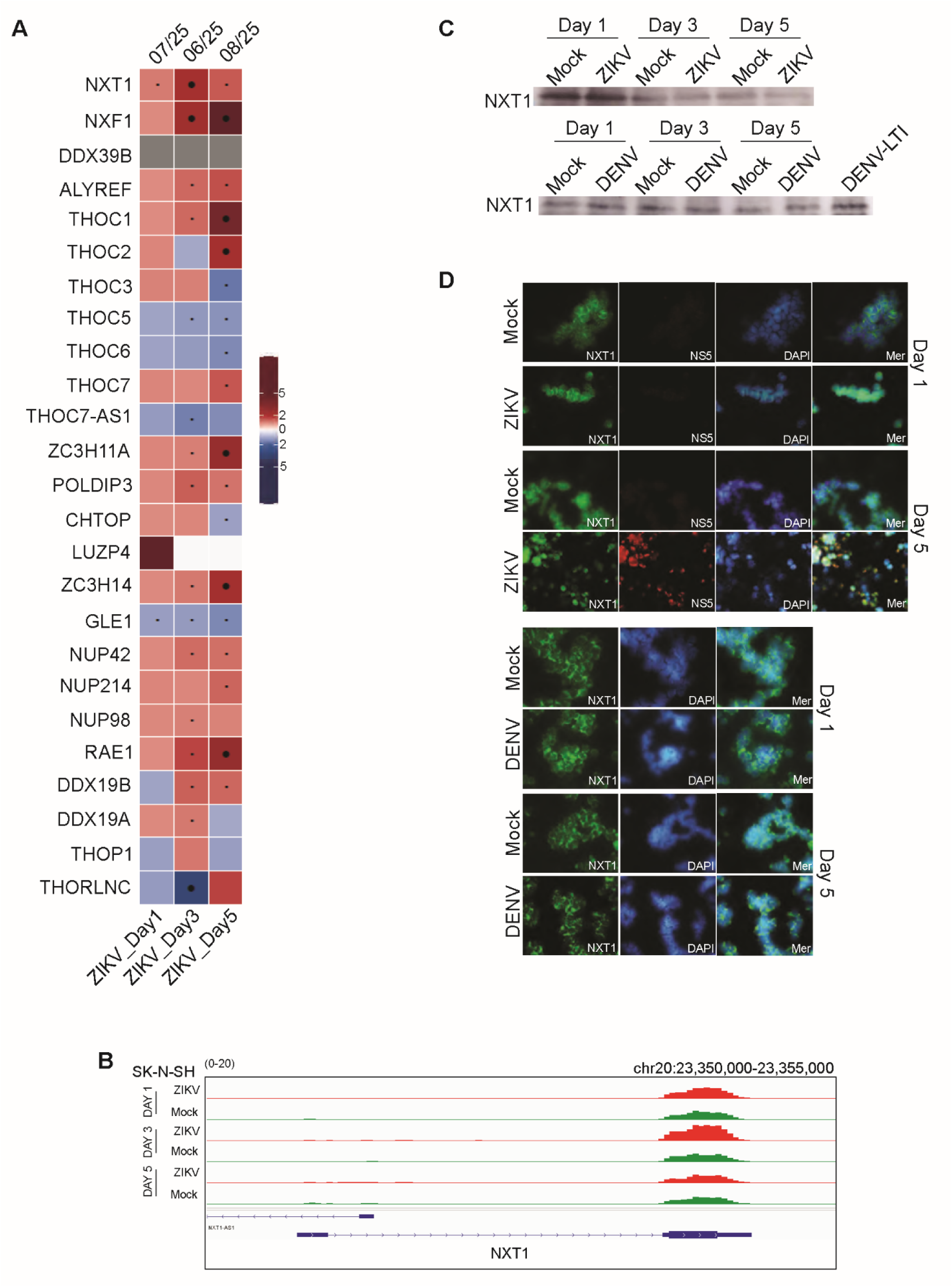
Comparison of NPC pathway transcripts in mock, ZIKV-infected SK-N-SH alongside protein level analysis. (A) Heatmap of NPC pathway-associated normalized RNA-seq transcripts for mock versus ZIKV-infected SK-N-SH samples at 1-, 3-, and 5-DPI. (B) A comparison of normalized RNA-seq reads represented in reads per million for NXT1 and visualized in IGV. Comparisons shown for mock and ZIKV-infected SK-N-SH samples at 1-, 3-, 5-DPI. (C) Western blot analysis of anti-NXT1 for mock, ZIKV-, and DENV-infected samples at 1-, 3-, 5-DPI as well as long-term DENV-infected sample. (D) Immunocytochemistry staining of mock, ZIKV, and DENV at 1- and 5-DPI. Antibodies used were anti-NXT1 (green), anti-ZIKV NS5 (red), and DAPI (blue).

**Fig. S12.**
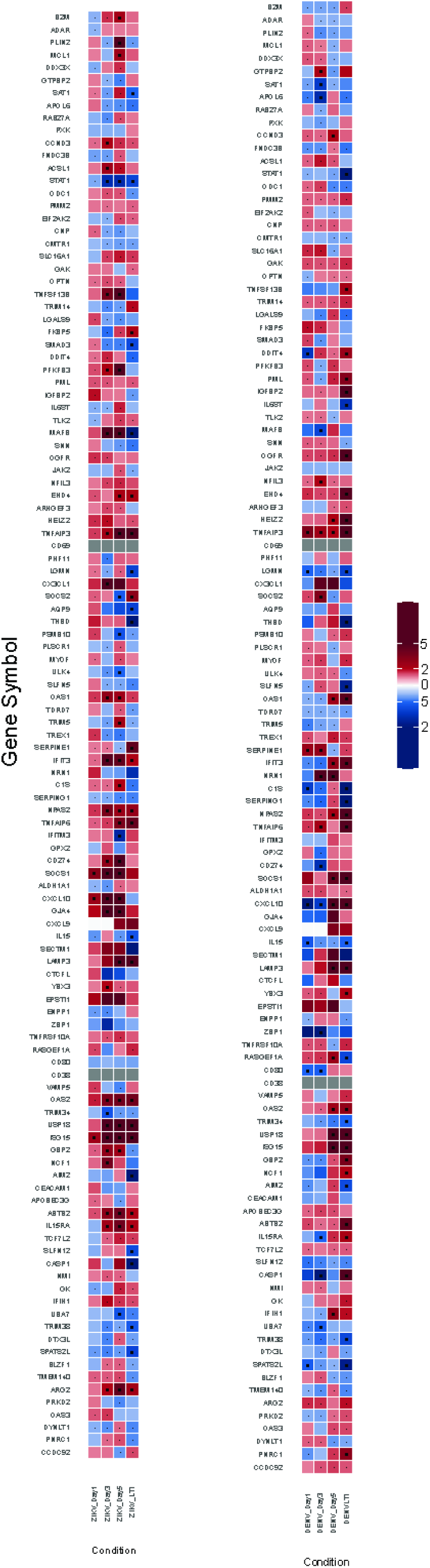
Comparison of NMD target transcripts in Vero acute and long-term infections. Heatmap of normalized RNA-seq data for mock versus ZIKV-infected (left) and DENV-infected (right) Vero samples at 1-, 3-, 5-DPI, and long-term infected samples.

